# Post-transcriptional regulatory pre-complex assembly drives timely cell-state transitions during differentiation

**DOI:** 10.1101/2024.04.29.591706

**Authors:** Hideyuki Komori, Geeta Rastogi, John Paul Bugay, Hua Luo, Sichun Lin, Stephane Angers, Craig A. Smibert, Howard D. Lipshitz, Cheng-Yu Lee

**Affiliations:** Life Sciences Institute, University of Michigan, Michigan, Ann Arbor 48109, USA; Department of Cell and Developmental Biology, University of Michigan Medical School, Ann Arbor 48109, USA; Division of Genetic Medicine, Department of Internal Medicine and Comprehensive Cancer Center, University of Michigan Medical School, Ann Arbor, 48109, USA; Department of Molecular Genetics, University of Toronto, Toronto, ON M5G 1M1, Canada; Donnelly Centre for Cellular and Biomolecular Research, University of Toronto, Toronto, ON M5S 3E1, Canada; Department of Biochemistry, University of Toronto, Toronto, ON M5G 1M1, Canada

## Abstract

Complexes that control mRNA stability and translation promote timely cell-state transitions during differentiation by ensuring appropriate expression patterns of key developmental regulators. The *Drosophila* RNA-binding protein Brain tumor (Brat) promotes degradation of target transcripts during the maternal-to-zygotic transition in syncytial embryos and in uncommitted intermediate neural progenitors (immature INPs). We identified Ubiquitin-specific protease 5 (Usp5) as a Brat interactor essential for the degradation of Brat target mRNAs in both cell types. Usp5 promotes Brat-dedadenylase pre-complex assembly in mitotic neural stem cells (neuroblasts) by bridging Brat and the scaffolding components of deadenylase complexes lacking their catalytic subunits. The adaptor protein Miranda binds the RNA-binding domain of Brat, limiting its ability to bind target mRNAs in mitotic neuroblasts. Cortical displacement of Miranda activates Brat-mediated mRNA decay in immature INPs. We propose that the assembly of an enzymatically inactive and RNA-binding-deficient pre-complex poises mRNA degradation machineries for rapid activation driving timely developmental transitions.

## Introduction

Timely cell-state transitions during differentiation are essential for normal development, and defects in these transitions can lead to birth disorders and death ^1-6^. Gene products that are no longer required or define a different cell-state are often cleared from the cell through mechanisms that act at the transcriptional and post-transcriptional levels to allow new deterministic gene products to function in a timely manner. Intensive efforts have provided significant insights into how transcriptional and post-translationally mechanisms downregulate protein expression to facilitate cellular differentaiton. By contrast, less is known about how the transcript degradation is precisely timed and executed during cell-state transitions.

The cell-type hierarchy of neuroblast lineages in the fly larval brain has been characterized at both the functional and molecular levels ^7-9^. A wild-type larval brain lobe contains approximately 100 neuroblasts each of which divides asymmetrically every 60-90 minutes to regenerate itself and to produce a sibling progeny that commits to differentiation. Eight of these neuroblasts are Type II and are characterized by the generation of an immature INP in every asymmetric division. Notch signaling must be robustly and rapidly terminated at all levels in newborn immature INPs to permit commitment to an INP identity, which occurs approximately 75 minutes after neuroblast division ^10-14^. The Brat RNA-binding protein (RBP) localizes to the basal cortex of mitotic Type II neuroblasts and segregates into newborn immature INPs where Brat promotes the decay of Notch downstream-effector gene transcripts ^14-20^. Loss of *brat* function results in aberrant translation of the Notch downstream-effector gene transcripts *deadpan* (*dpn*), *klumpfuss* (*klu*) and *zelda* in newborn immature INPs, driving their reversion to supernumerary Type II neuroblasts ^14,21-24^. Numb, a conserved antagonist of Notch signaling, also asymmetrically segregates to newborn immature INPs ^17,19,25-27^. Loss of *numb* function leads to ectopic Notch signaling and transcription of *dpn*, *klu* and *zelda* in newborn immature INPs. The increased levels of *dpn*, *klu* and *zelda* transcripts overwhelm the Brat mRNA decay machinery in *numb*-mutant immature INPs leading to their ectopic translation and the formation of supernumerary Type II neuroblasts. The severity of this supernumerary neuroblast phenotype is directly proportional to the strength of the *numb* mutant alleles ^28^. Thus, the Brat-regulated cell-state transition in newborn immature INPs provides a highly sensitive platform for investigating at single-cell resolution the assembly and activation of RBP-based complexes that act to destabilize transcripts that would otherwise disrupt cell differentiation.

Brat is a member of the conserved TRIM-NHL family of RBPs and is characterized by two N-terminal B-boxes, a central coiled-coil domain, and a C-terminal NHL domain ^1^. Brat functions through the B-boxes to recruit proteins that are essential for Brat-mediated mRNA decay in newborn immature INPs while binding the Brat-responsive element (BRE; UGUUA) in the 3’UTRs of target transcripts via the NHL domain ^23,29-31^. Tis11 (the fly homolog of vertebrate Tristetraprolin) interacts with Brat via the B-boxes, and functions together with Brat to degrade Notch downstream-effector gene transcripts in newborn immature INPs ^14^. Tis11 recognizes the AUUUA pentamer sequence (AU-rich element or ARE) in the 3’UTRs of target transcripts and promotes their decay by recruiting the CCR4-NOT deadenylase complex ^32-34^. Although high-affinity BREs are frequently located in tandem with AREs, overexpressing RNA-binding-defective Tis11 can partially substitute wild-type Tis11 in Brat-mediated repression of Notch downstream-effector gene transcripts in newborn immature INPs ^14^. Thus, Tis11 likely promotes robust Brat-mediated mRNA decay.

We investigated the control of Brat-mediated mRNA decay during the transition from a neuroblast state to the onset of INP commitment in newborn immature INPs. Immunoprecipitation of Brat followed by mass spectrometry (IP-MS) identified Ubiquitin-specific protease 5 (Usp5) as an RNA-independent Brat binding partner. We show that Usp5 functions through the conserved Cysteine residue in the enzymatic domain to asymmetrically localize and segregate into immature INPs via a Brat-dependent mechanism. Usp5 is required for timely clearance of Brat target mRNAs in early embryos as well as in newborn immature INPs in larval brains. Usp5 promotes Brat-mediated co-localization and co-segregation of scaffolding components of the CCR4-Not1 and Pan2-Pan3 deadenylase complexes during asymmetric neuroblast division. Because the enzymatic components of both deadenylase complexes do not asymmetrically localize in mitotic neuroblasts, we propose that Usp5 functions as a scaffold to promote the assembly of a Brat-deadenylase pre-complex that is enzymatically inactive. The adopter protein Miranda (Mira) binds the coiled-coil and NHL domains of Brat and asymmetrically localizes Brat in mitotic neuroblasts ^18,23^. Because the NHL domain also mediates Brat-mRNA interactions ^18,23^, Mira-mediated sequestration of Brat limits its RNA binding capacity. Mira is displaced from the cortex of newborn immature INPs and releases its cargo proteins ^35,36^. Mutant Mira that constitutively associates with the cortex maintains the Brat pre-complex leading to aberrant translation of Brat target transcripts driving their reversion to supernumerary neuroblasts. Thus, cortical release of Mira promotes activation of the Brat-deadenylase pre-complex in newborn immature INPs, allowing activation of the Brat mRNA decay complex. We propose that assembling an enzymatically inactive and RNA-binding deficient pre-complex poises the mRNA decay machineries for rapid activation during cell-state transitions, and will likely be relevant in most, if not all, similar biological processes throughout metazoans.

## Results

### Usp5 interacts with Brat and promotes timely clearance of Brat target transcripts during the MZT

Because Brat is required for the degradation of maternally deposited transcripts in the early embryo, we searched for candidate Brat-interacting proteins that contribute to Brat-mediated mRNA decay in early embryos. We immunoprecipitated Brat-interacting proteins using untreated or RNase-treated lysate extracted from 0-3-hr old embryos (Fig. 1A) and determined the identities of Brat-interacting proteins by liquid chromatography-tandem mass spectrometry (LC-MS/MS). We defined Brat interactors as proteins that were enriched in the Brat immunoprecipitate with significant analysis interactome (SAINT ^37^) score of ≥ 0.9 and Bayesian false discovery rate (BFDR) of < 0.05. Supernumerary limbs (Slmb), Ubl-specific protease (Ulp1) and Ubiquitin specific protease 5 (Usp5) were present in both Brat immunoprecipitates suggesting that these proteins directly interact with Brat (Fig. S1A and Table S1). We excluded Slmb, a conserved F-box protein and an essential component of several E3 ubiquitin-ligase complexes, from further consideration because these complexes degrade a wide variety of substrates including proteins and mRNAs ^38,39^. We focused on Usp5 instead of Ulp1 in this study due to the availability of reagents.

**Figure 1.**
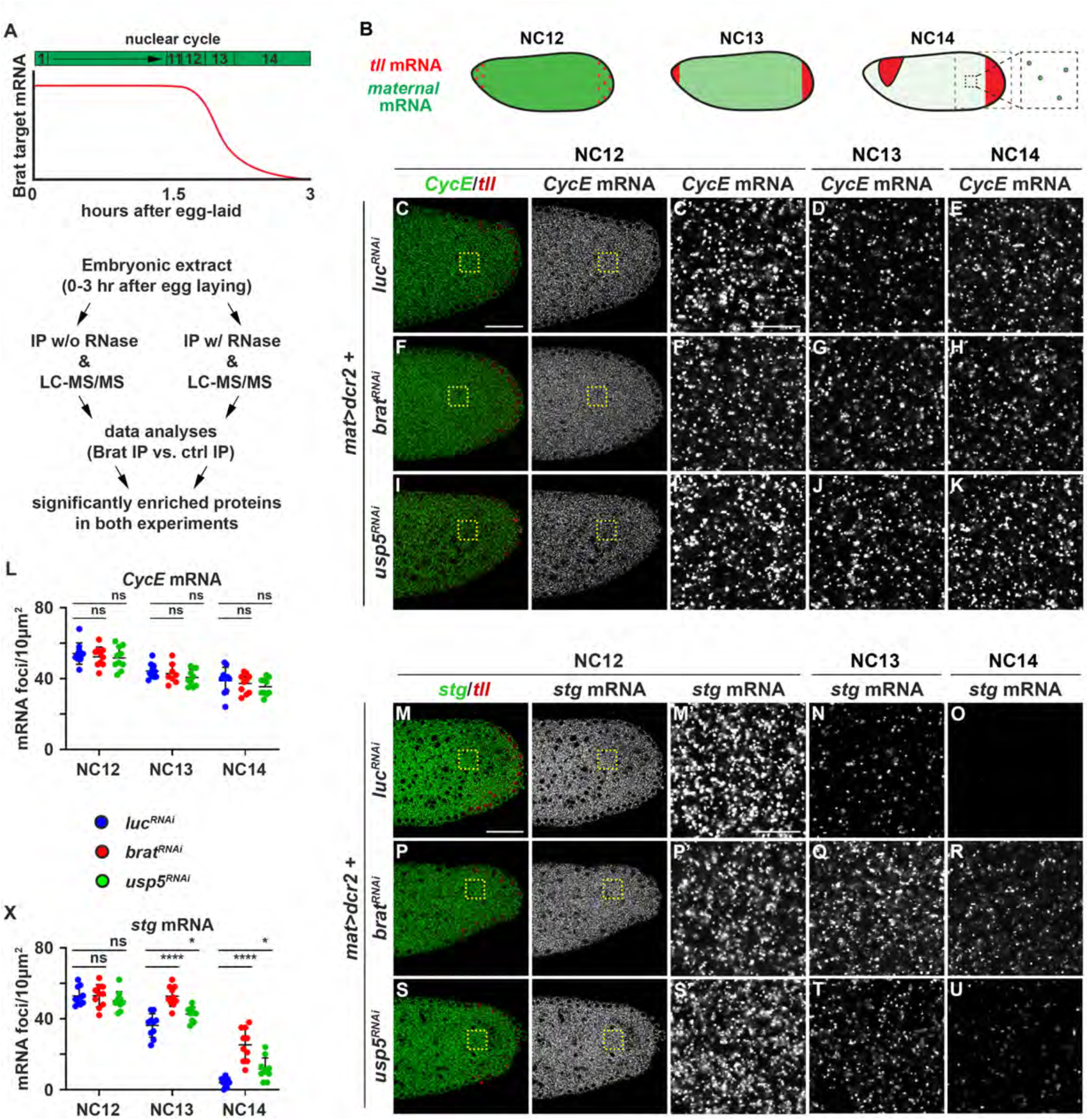
Usp5 interacts with Brat and promotes timely clearance of Brat target mRNAs during MZT. **A,** Top: Schematic showing the timing of the clearance of maternally deposited Brat-target mRNA decay during the MZT. Y-axis: the level of Brat target mRNAs. X-axis: hours after egg deposition. Bottom: The IP-MS strategy to identify Brat-interacting proteins using extracts from 0-3 hour embryos. **B,** Degradation of maternally deposited mRNAs and expression pattern of zygotically transcribed *tll* mRNAs in NC12, NC13, and NC14. The large square marks the posterior of the embryo where low-magnification smFISH images were taken. The small square represents the 10μm^2^ area anterior to zygotic *tll* mRNAs where high-magnification smFISH images were taken. **C-K,** NC12-NC14 stage embryos with *luciferase (luc)* (C-E), *brat* (F-H) or *usp5* (I-K) maternal knock-downs were co-hybridized with smFISH probes for *CycE* and *tll*. **L,** Quantification of *CycE* mRNA foci in the 10μm^2^ area shown in the hi-magnification images. **M-U,** NC12-NC14 stage embryos with *luc* (M-O), *brat* (P-R), and *usp5* (S-U) maternal knock-down were co-hybridized with smFISH probes for *stg* and *tll*. **X,** Quantification of *stg* mRNA foci in the 10μm^2^ area shown in hi-magnification images. The following labels apply to all images in this figure. Yellow dashed squares indicate the location of the high magnification images. Scale bars in low-magnification images: 50μm. Scale bars in hi-magnification images: 10μm. Graphs show mean number ± standard deviation. P-values: *<0.05, **<0.01, ***<0.001, ****<0.0001. “ns” indicates no significance.

If Usp5 functions together with Brat to promote degradation of maternally deposited mRNAs in the embryo, transcripts that depend on Brat for clearance should ectopically persist in *usp5*-mutants (Fig. 1A). The *brat* and *usp5* transcripts are maternally deposited in oocytes (Fig. S1B,D). Overexpressing a *UAS-brat^RNAi^*or *UAS-usp5^RNAi^* transgene driven by a maternal *α-Tub-Gal4* driver drastically reduced their transcript levels allowing for the generation of *brat* or *usp5* ‘maternal-mutant’ embryos (Fig. S1C,E). Brat promotes decay of maternally deposited transcripts during nuclear cycle (NC) 12-14, which can be unambiguously defined by the localization patterns of zygotic *tailless* (*tll*) transcripts ^2,40^ (Fig. 1B). We quantified the number of non-Brat target – *Cyclin E* (*CycE*) and *mRpL15* – or Brat target mRNA foci by single-molecule fluorescent *in situ* hybridization (smFISH) in a 10μm^2^ area anterior to the *tll* domain. Overall levels of non-Brat target mRNA foci progressively declined from NC12 to NC14 in control (*luciferase_RNAi_*), *brat* and *usp5* maternal-mutant embryos (Fig. 1C-L; Fig. S1F). Importantly, levels of non-Brat target mRNAs were statistically indistinguishable between the three maternal-mutant conditions in each of the three stages. Brat target mRNA levels – *string* (*stg*), *treholase* (*treh*), *hunchback* (*hb*) and *Glutamate Carrier 1* (*GC1*) – were indistinguishable between control and mutant embryos in NC12 but were significantly elevated in NC13 and 14 in *brat* or *usp5*-maternal mutant embryos relative to control embryos (Fig. 1M-X; Fig. S1G-I). Together these results indicate that Usp5 is a Brat interactor and is required for timely clearance of Brat target mRNAs during the MZT.

### Usp5 depends on Brat for asymmetric localization in mitotic neuroblasts and segregation into INPs

We tested whether Usp5 might be a Brat interactor in neuroblasts by first determining its localization and segregation pattern during asymmetric neuroblast division in wild-type brains. The adapter protein Mira asymmetrically localizes Brat in the basal cortex of metaphase neuroblasts and segregates Brat into the cortex of future newborn immature INP in telophase (Fig. 2A) ^18,23^. Thus, Mira localization patterns serve as a reliable readout for the spatial distribution of Brat in mitotic neuroblasts. We found that Usp5 co-localized with Mira in a basal crescent in metaphase neuroblasts and co-segregated with Mira into the cortex of immature INPs at telophase in wild-type brains (Fig. 2B,C). If Brat-binding is required for asymmetric localization and segregation of Usp5 in mitotic neuroblasts, loss of *brat* function should lead to symmetric segregation of Usp5. Asymmetric localization of Mira is not dependent on Brat and served as a control in this experiment. Usp5 was uniformly distributed in the cytoplasm of metaphase neuroblasts and segregated into both neuroblast progeny in *brat*^-/-^ brains (Fig. 2D,E). Importantly, Brat remained basally localized in mitotic neuroblasts and segregated into the cortex of future newborn immature INP in *usp5*^-/-^ brains (Fig. 2F,G). These data indicate that Brat is required for asymmetric localization in mitotic neuroblasts and segregation of Usp5 into INPs, supporting Usp5 as a Brat interactor during asymmetric neuroblast division.

**Figure 2.**
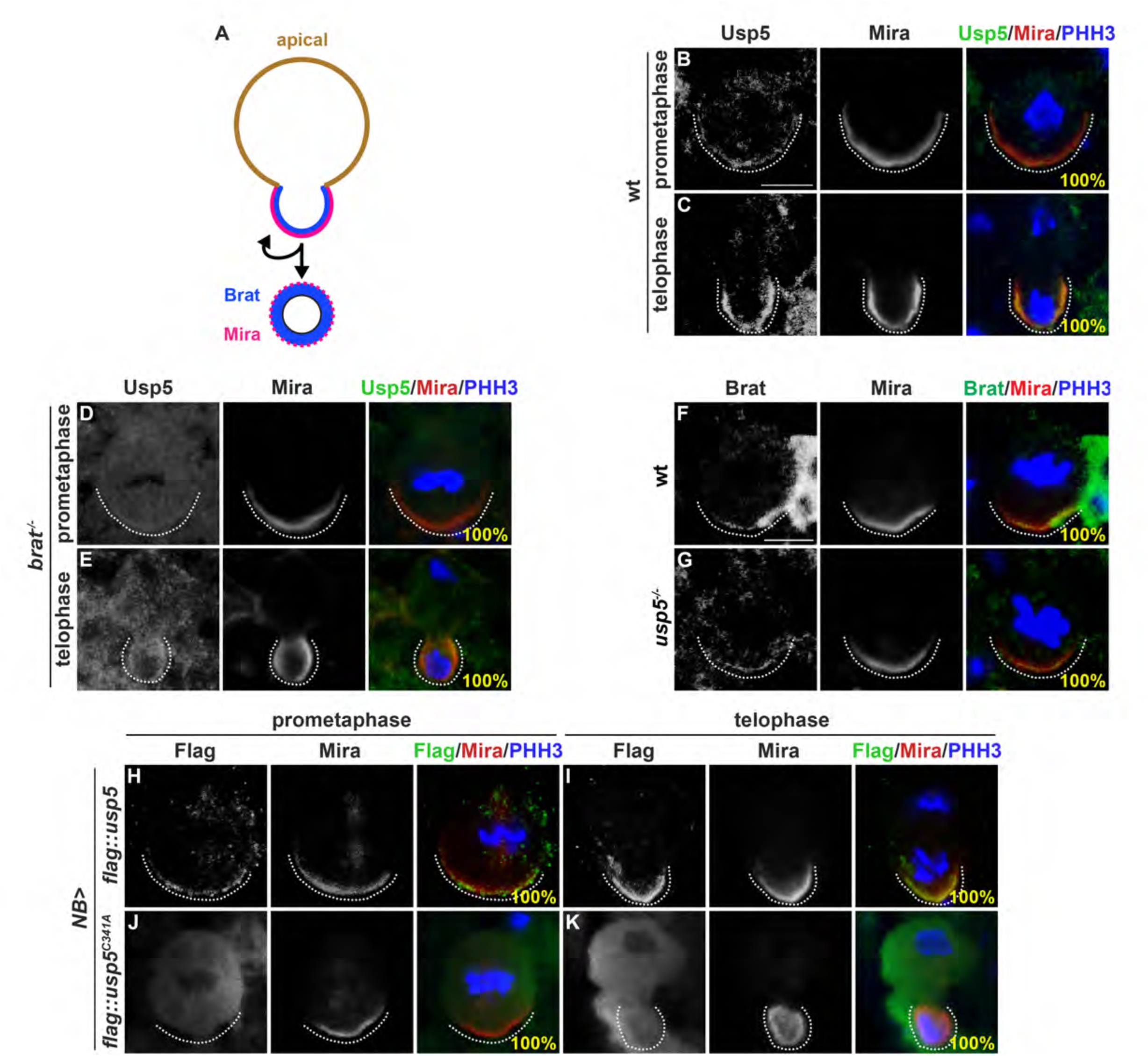
Usp5 depends on Brat for basal localization and segregation during asymmetric neuroblast division. **A,** A schematic showing the localization pattern of Mira (pink) and Brat (blue) during asymmetric neuroblast division. Dashed line indicates Mira displaced from the cortex of newborn immature INP. **B-G,** Mitotic neuroblasts of the indicated genotypes were co-stained with antibodies against the indicated proteins. The localization pattern of Usp5 in prometaphase or telophase neuroblasts in wild-type (B-C) or *brat^-/-^* (D-E) brains. The localization pattern of Brat in prometaphase or telophase neuroblasts in *usp5^-/-^* (F-G) brains. **H-K,** Mitotic neuroblasts that overexpress either a FLAG-tagged wild-type Usp5 or Usp5^C341A^ were co-stained with specific antibodies against FLAG, Mira and Phosphohistone H3 (PHH3). For all images in this figure: Scale bars, 5μm. White dotted lines outline the basal cortex of neuroblasts. wt: wild-type. *brat^-/-^*: *brat^11^/Df(2L)Exel8040. usp5^-/-^*: *usp5^leon1/leon1^*.

Usp5 is a member of the Ubiquitin Specific Protease (USP) family of deubiquitinases, and the Cysteine residue in the ubiquitin-specific processing protease domain is conserved in all USPs from yeast to human ^41^. We tested if the conserved Cysteine residue (C341 in fly Usp5) is required for asymmetric localization and segregation of Usp5 in mitotic neuroblasts. Overexpressed Usp5::Flag asymmetrically localized and segregated during asymmetric neuroblast division (Fig. 2H,I). By contrast, overexpressed Usp5^C341A^::Flag was distributed throughout the cytoplasm of metaphase neuroblasts and both neuroblast progeny (Fig. 2J,K). Thus, C341 is required for Brat-mediated polarized localization and segregation of Usp5 during asymmetric neuroblast division and may facilitate Usp5 interaction with Brat in neuroblasts.

### Usp5 functions together with Brat to promote INP commitment in immature INPs

We next tested if Usp5 promotes the onset of INP commitment in newborn immature INPs. First, we quantified total Type II neuroblasts (Dpn^+^Ase^-^ cells with ≥ 9 μm in diameter) per brain lobe in age-matched wild-type versus *usp5*-null larvae. Two *usp5*-mutant alleles were previously reported ^42^. *usp5^leon1^*is a strong loss-of-function allele and *usp5^leon2^* is a hypomorphic allele (Fig. 3A). We generated a *usp5*-null allelic combination by crossing *usp5^leon1^* to a *usp5^Df^* (*leon^19-2^*) that removes the *usp5* locus. *usp5*-null larvae died in the third larval instar and were significantly smaller than identically staged control larvae. *usp5*-null brain lobes contained statistically indistinguishable numbers of Type II neuroblasts relative to a wild-type brain lobes (8.4 ± 1 vs. 8.0 ± 0; p = 0.3) (Fig. 3B-D). Unlike wild-type brains, Ase^-^ immature INPs in *usp5*-null brains expressed Dpn, suggesting defects in down-regulation of Notch downstream-effector gene expression (Fig. 3C). Because Usp5 is required for cell viability ^43^, loss of cell viability could mask the reversion of Ase^-^ immature INPs to supernumerary neuroblasts in *usp5-null* brains. Mis-expression of baculovirus apoptotic inhibitor p35 in wild-type brains did not result in supernumerary neuroblast formation (8.0 ± 0 Type II neuroblasts per lobe). By contrast, mis-expression of p35 in *usp5-null* brains led to supernumerary neuroblast formation (14.9 ± 3.5 Type II neuroblasts per lobe) (Fig. 3D,E). Importantly, overexpressing wild-type Usp5::Flag but not Usp5^C341A^::Flag rescued the supernumerary neuroblast phenotype in *usp5-null* brains mis-expressing p35 (Fig. 3D,F,G). These data indicate that Usp5 functions through C341 to promote INP commitment in immature INPs.

**Figure 3.**
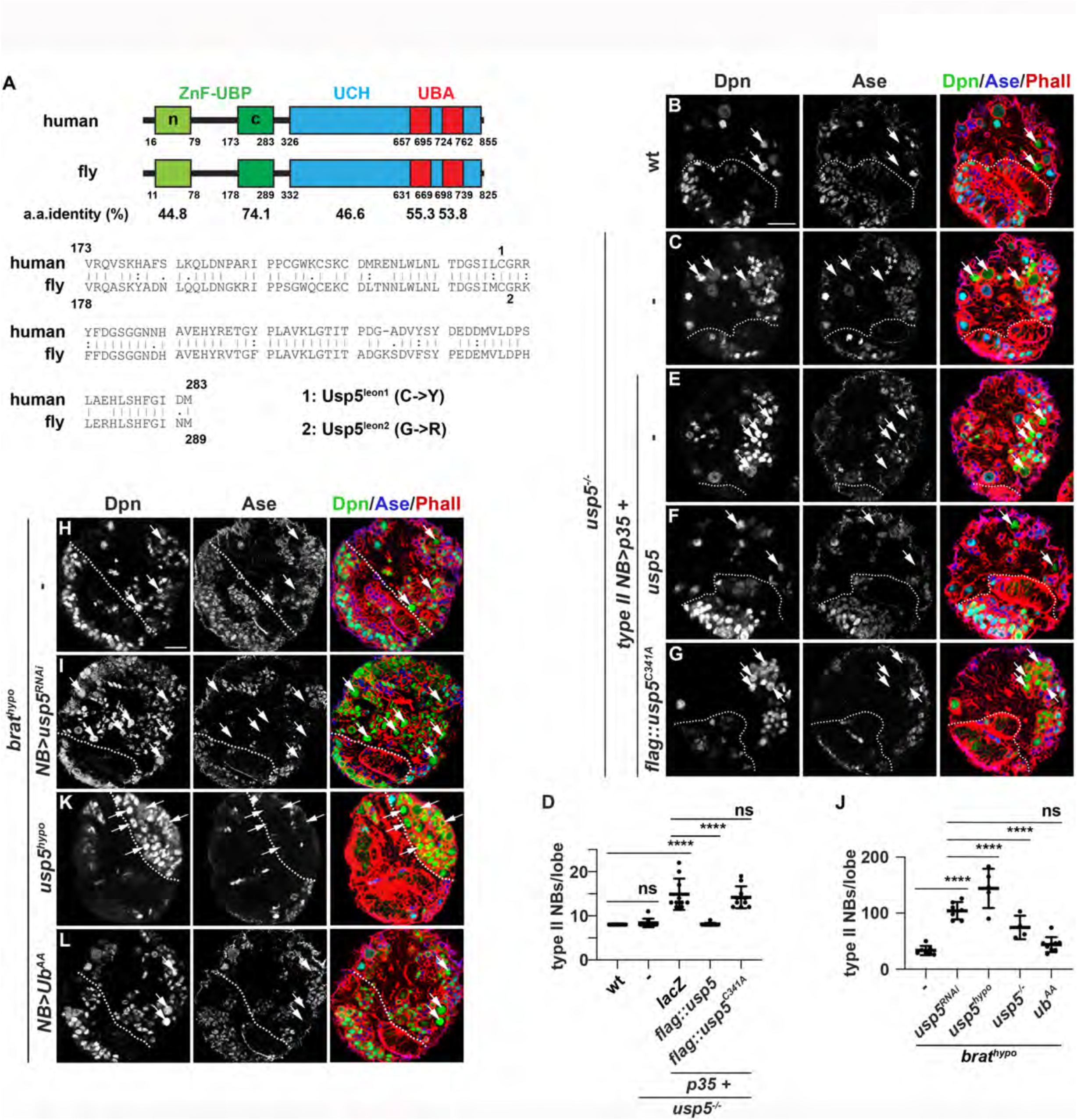
Usp5 functions in Brat-mediated mRNA decay in newborn immature INPs independent of its role in regulating ubiquitin homeostasis. **A,** The protein domain structure of human and fly Usp5 and the amino acid substitution in two previously isolated *usp5* alleles. ZnF-UBP: zinc-finger ubiquitin-binding domain, UCH: Ubiquitin carboxyl-terminal hydrolase, UBA: Ubiquitin-associated domain. **B-G,** *usp5*^-/-^ brains alone or overexpressing the indicated *UAS* transgenes co-stained with specific antibodies against Dpn, Ase and Phalloidin. (D) Quantification of total Type II neuroblasts per brain lobe of the indicated genotype. n = 10 brains per genotype. **H-L,** *brat^hypo^*brains alone or overexpressing indicated *UAS* transgenes were co-stained with specific antibodies against Dpn, Ase and Phalloidin. (J) Quantification of total Type II neuroblasts per brain lobe of the indicated genotype. n = 4-10 brains per genotype. “-“ indicates no transgene. Scale bars, 20μm. White dotted line separates the optic lobe from the central brain. White arrows: Type II neuroblast (Dpn^+^Ase^-^; <u>≥</u> 9 μm in diameter). White asterisk: Ase^-^ immature INPs aberrantly expressing Dpn (<u>≤</u> 5 μm in diameter). wt: wild-type. *brat^hypo^*: *brat^DG19310/11^*. *usp5^-/-^*: *usp5^leon1leon1^*. *usp5^hypo^*: *usp5^leon1^/^leon2^*. *usp5^RNAi^*: *TRiP.JF02163*. *NB>*: *Wor-Gal4*. *Type II NB>*: *Wor-Gal4,Ase-Gal80*. Graphs show mean ± standard deviation. P-values: ***<0.001, ****<0.0001. “ns” indicates no significance.

Usp5 functions in ubiquitin homeostatis, and as such we explored the role of Usp5-mediated ubiquitin regulation in Brat function. N-terminus of Usp5 contains two zinc-finger ubiquitin-binding domains (nZn-UBP and cZn-UBP) (Fig. 3A) ^44,45^. nZn-UBP is required for the deubiquitinase activity of Usp5 to hydrolyze the di-glycine tail in unanchored polyubiquitin chains into free monoubiquitin ^46,47^. By contrast, cZn-UBP is largely dispensable for the catalytic activity of Usp5 but, rather, promotes protein-protein interactions including binding to ubiquitin. The *usp5^leon1^* and *usp5^leon2^* mutations lead to a single amino acid substitution at Cysteine 224 or Glycine 225 in the cZn-UBP and drastic expansion of postsynaptic specializations of neuromuscular junctions (Fig. 3A) ^42,43^. Overexpressing Ub^AA^, which carries amino acid substitutions at Gly75Ala and Gly76Ala and cannot be cleaved by Usp5-mediated proteolysis, mimics the *usp5^leon1^* or *usp5^leon2^* phenotype at neuromuscular junctions ^43^. These observations suggest that the *usp5^leon1^* or *usp5^leon2^* mutation perturbs Usp5 interaction with unanchored poly-Ubiquitin chains leading to defects in ubiquitin homeostasis. We, therefore, tested whether Usp5 promotes INP commitment in immature INPs by regulating ubiquitin homeostasis by overexpressing Ub^AA^ in a *brat*-hypomorphic (*brat*^hypo^) genetic background that is highly sensitive to changes in Brat-mediated mRNA decay ^14^. Consistent with Usp5 functioning together with Brat in a complex, knocking down *usp5* function by *RNAi* drastically enhanced the supernumerary Type II neuroblast phenotype in *brat*^hypo^ brains (Fig. 3H-J). The *usp5*-hypomorphic (*usp5^leon1^*/*^leon2^*; *usp5^hypo^*) or *usp5*-null allelic combination also enhanced the supernumerary Type II neuroblast phenotype in *brat*^hypo^ brains (Fig. 3J,K). Ub^AA^ overexpression led to ectopic accumulation of polyubiquitin in Type II neuroblasts but did not lead to supernumerary neuroblast formation in wild-type brains (Fig. S2). Most importantly, Ub^AA^ overexpression did not significantly enhance the supernumerary Type II neuroblast phenotype in *brat^hypo^*brains (Fig. 3J,L). These data suggest that Usp5 is unlikely to promote INP commitment in immature INPs by regulating ubiquitin homeostasis. Together, our analyses of Usp5 localization patterns and *usp5*-mutant phenotypes during asymmetric neuroblast division support a model in which Usp5 functions together with Brat to promote INP commitment independent of its enzymatic activity.

### Usp5 promotes clearance of Brat target transcripts in immature INPs

The Notch downstream-effector genes *dpn* and *E(spl)mγ* and cell cycle genes such as *CycE* are highly expressed in Type II neuroblasts but become rapidly downregulated in newborn immature INPs (Fig. S3A). Downregulation of *dpn* depends on Brat-mediated mRNA decay but *E(spl)mγ* and *CycE* do not, indicating that Dpn^+^ cells in *brat*-null brains may be either supernumerary neuroblasts or immature INPs aberrantly expressing Dpn due to loss of *brat* function. To test whether Usp5 functions in Brat-mediated mRNA decay, we first had to establish unambiguous cell identity markers unaffected by loss of Brat-mediated mRNA decay in the Type II neuroblast lineage and a strategy for quantifying mRNA levels. smFISH detected high levels of *dpn* mRNAs in Type II neuroblasts (Dpn^+^,Erm::V5^-^; >9μm in diameter) and INPs (Dpn^+^,Erm::V5^-^; <5μm in diameter), drastically lower levels in newborn immature INPs (Dpn^-^,Erm::V5^-^), and virtually undetectable levels in immature INPs (Dpn^-^,Erm::V5^+^) in the wild-type Type II neuroblast lineage (Fig. 4A,B). *CycE* mRNAs displayed an expression pattern identical to *dpn* and *E(spl)mγ* gene products (both transcripts and protein) in the wild-type Type II neuroblast lineage (Fig. 4C,D; Fig. S3A). We combined *dpn*, *E(spl)mγ* and *CycE* expression to define the identities of cells in 24- or 48-hr RFP-marked mosaic clones derived from single *brat*-null Type II neuroblasts. While Dpn was detected in all cells in the 24-hr clones, E(spl)mγ::GFP was only detectable in parental neuroblasts (>9μm in cell diameter) (Fig. S3B). Dpn and E(spl)mγ::GFP were co-expressed in parental neuroblasts and several supernumerary neuroblasts (5-9μm in cell diameter) in 48-hr clones while all remaining cells in the clones expressed only Dpn (Fig. S3C). These results suggest that most cells in *brat*-null Type II neuroblast clones are immature INPs that aberrantly express Dpn due to loss of Brat-mediated decay of *dpn* transcripts. Consistently, 1-2 supernumerary neuroblasts (E(spl)mγ::GFP^+^ and *CycE* mRNA*^+^*) were first detected in 36-hour *brat-*null clones (Fig. S3D-H). These data indicate that all neuroblast progeny in *brat*-null Type II neuroblast clones aged for 30 hours or less are immature INPs and should display ectopic Brat target mRNAs.

**Figure 4:**
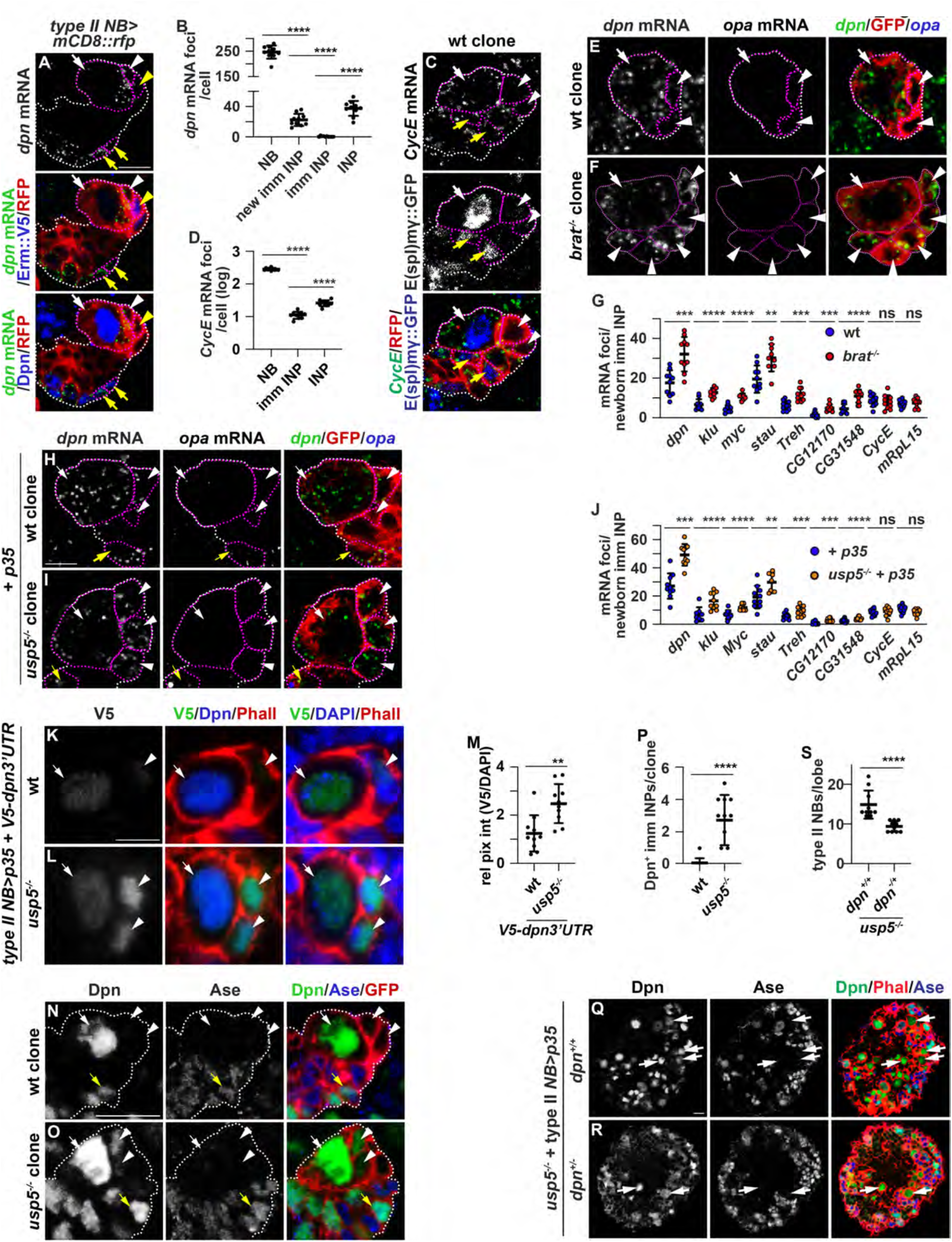
Usp5 promotes timely clearance of Brat target transcripts in immature INPs. **A,** *erm::V5* third-instar larval brains overexpressing mCD8::RFP marking Type II neuroblast lineages were hybridized with smFISH probes for *dpn* and counter-stained with antibodies against V5 and RFP. Scale bars, 10μm. **B,** Quantification of *dpn* mRNA foci per indicated cell types per brain lobe. n ≥ 10 brains. **C,** Larval brains carrying a *E(spl)mγ::GFP* transgene and containing RFP-marked wild-type Type II neuroblast mosaic clones aged for 96 hours following induction were hybridized with smFISH probes for *CycE* and counter-stained with specific antibodies against GFP and RFP. Scale bars, 5μm. **D,** Quantification of *CycE* mRNA foci per indicated cell type per brain lobe. n ≥ 10 brains. **E-G,** Larval brains containing GFP-marked wild-type or *brat*^-/-^ Type II neuroblast mosaic clones aged for 24 hours following induction were co-hybridized with smFISH probes for the indicated genes and counterstained with an antibody against GFP. Scale bars, 5μm. (G) Quantification of indicated mRNA foci in newborn immature INP in wild-type and *brat^-/-^*immature INPs (absence of *opa* transcripts). n = 8-10 clones. **H-J,** Larval brains containing GFP-marked wild-type or *usp5*^-/-^ Type II neuroblast mosaic clones overexpressing P35 and aged for 96 hours following induction were co-hybridized with smFISH probes for the indicated genes and counterstained with an antibody against GFP. Scale bars, 5μm. (J) Quantification of indicated mRNA foci in newborn immature INPs in wild-type and *usp5^-/-^* clones (absence of *opa* transcripts). n = 8-10 clones. **K-M,** wild-type or *usp5*^-/-^ larval brains carrying a *V5-dpn3’UTR* reporter transgene and overexpressing p35 were stained with antibodies against V5, Dpn, DAPI and Phalloidin. Scale bars, 5μm. (M) Quantification of relative reporter expression in newborn immature INPs of the indicated genotypes. The relative reporter expression levels were determined by calculating the ratio of V5 to DAPI. n = 10 brains. **N-P,** Larval brains containing GFP-marked wild-type or *usp5*^-/-^ Type II neuroblast mosaic clones aged for 96 hours following induction were stained with antibodies against Dpn, Ase and GFP. Scale bars, 5μm. (P) Quantification of Dpn-expressing immature INPs per brain lobe of the indicated genotype. n = 10 brains. **Q-S,** *usp5*^-/-^ larval brains overexpressing P35 alone or heterozygous for *dpn* were stained with specific antibodies against Dpn, Ase and Phal. Scale bars, 20μm. (S) Quantification of total Type II neuroblasts per brain lobe of the indicated genotypes. n = 10. *Type II NB>*: *Wor-Gal4,Ase-Gal80*. White dotted lines outline Type II neuroblast clones. Magenta dotted lines outline the cell boundary. White arrows: Type II neuroblast. White arrowheads: newborn immature INP. Yellow arrows: INP. Graphs show mean ± standard deviation. P-values: *<0.05, **<0.01, ***<0.001, ****<0.0001. “ns” indicates no significance. wt: wild-type, *brat^-/-^*: *brat^11/11^, usp5^-/-^*: *usp5^leon1/leon1^*.

We examined the distribution of other Brat target mRNAs in 24-hour RFP-marked *brat-null* Type II neuroblast clones. *odd-paired* (*opa*) transcripts positively mark INPs, and *CycE* and *mRpL15* transcripts serve as negative controls due to the absence of BREs in their 3’UTRs ^14,28,48,49^. The number of control mRNA foci in Type II neuroblasts and immature INPs were statistically indistinguishable between wild-type and *brat*-null clones (Fig. 4E-G; Fig. S3I-K). While the number of Brat target mRNA foci in Type II neuroblasts remained statistically indistinguishable between these two genotypes, their abundance in immature INPs was significantly higher in *brat*-null clones than in wild-type clones (Fig. 4E-G; Fig. S3I-K). Thus, smFISH provides a reliable strategy for quantifying Brat target mRNA levels in Type II neuroblasts and immature INPs.

We next quantified Brat target transcripts in immature INPs in wild-type or *usp5*-null Type II neuroblast clones that overexpressed a *UAS-p35* transgene. The number of control mRNA foci in Type II neuroblasts and in immature INPs were indistinguishable between wild-type clones and those overexpressing p35, indicating that inhibiting apoptosis did not affect mRNA decay (Fig. 4G,J; Fig. S3K,N). The abundance of Brat target transcripts in Type II neuroblasts was statistically indistinguishable between wild-type and *usp5*-null clones overexpressing p35, but the number of Brat target mRNA foci in immature INPs was significantly higher in *usp5*-null brains (Fig. 4H-J; Fig. S3L-N). These data indicate that Usp5 is required for the decay of Brat target transcripts in immature INPs and support the hypothesis that Usp5 is a key component of the Brat mRNA decay complex.

To confirm the specificity of a role for Usp5 in Brat-mediated mRNA decay, we tested whether loss of *usp5* function aberrantly upregulates reporter activity controlled by the endogenous *dpn* 3’UTR or a synthetic Brat-responsive 3’UTR containing previously verified BREs ^14^. Relative expression levels of these reporters in Type II neuroblasts were indistinguishable between wild-type and *usp5*-null brains overexpressing p35 but were significantly higher in immature INPs in *usp5*-null brains than in wild-type brains (Fig. 4K-M; Fig. S3O-Q). Significantly more immature INPs in *usp5*-null clones ectopically expressed Dpn than in wild-type clones (Fig. 4N-P). Removing one copy of the *dpn* gene was sufficient to suppress the supernumerary neuroblast phenotypes in *usp5*-null brains (Fig. 4Q-S). These data support our model that Usp5 functions in the Brat complex to promote the transition from a neuroblast state to the onset of INP commitment by degrading Brat target mRNAs in immature INPs.

### Usp5 promotes assembly of the Brat pre-complex in mitotic neuroblasts

Loss of *usp5* function leads to ectopic translation of Brat target gene transcripts in immature INPs suggesting that Usp5-binding to Brat confers robust mRNA decay activity of the Brat complex. Because deadenylation of transcripts is the first step in mRNA decay ^50,51^, we tested whether Usp5 promotes the recruitment of the Pan2-Pan3 and CCR4-Not1 deadenylases to the Brat complex. We first focused on the scaffolding components of these complexes, Pan3 and Not1, and examined their localization patterns in mitotic neuroblasts in wild-type brains. We found that Pan3 and Not1 co-localized with Mira at a basal crescent in metaphase neuroblasts and asymmetrically segregated into the cortex of the future immature INP in telophase neuroblasts (Fig. 5A,B). By contrast, the enzymatic components of these complexes, Pan2 and Pop2, were uniformly distributed in the cytoplasm of mitotic neuroblasts and both progeny cells (Fig. 5C,D). These results suggest a model that scaffolding components of Pan2-Pan3 and CCR4-Not1 deadenylases might be in a complex with Brat and Usp5 in mitotic neuroblasts. Consistently, Pan3 and Not1 were uniformly distributed in the cytoplasm of mitotic neuroblasts and both neuroblast progeny in *brat*-null brains (Fig. 5E,F). Furthermore, Not1 also failed to asymmetrically localize and segregate in mitotic neuroblasts in *usp5*-null brains (Fig. 5G). These data suggest that Brat-Usp5-Not1-Pan3 form an enzymatically inactive pre-complex in mitotic neuroblasts (Fig. 5H).

**Figure 5.**
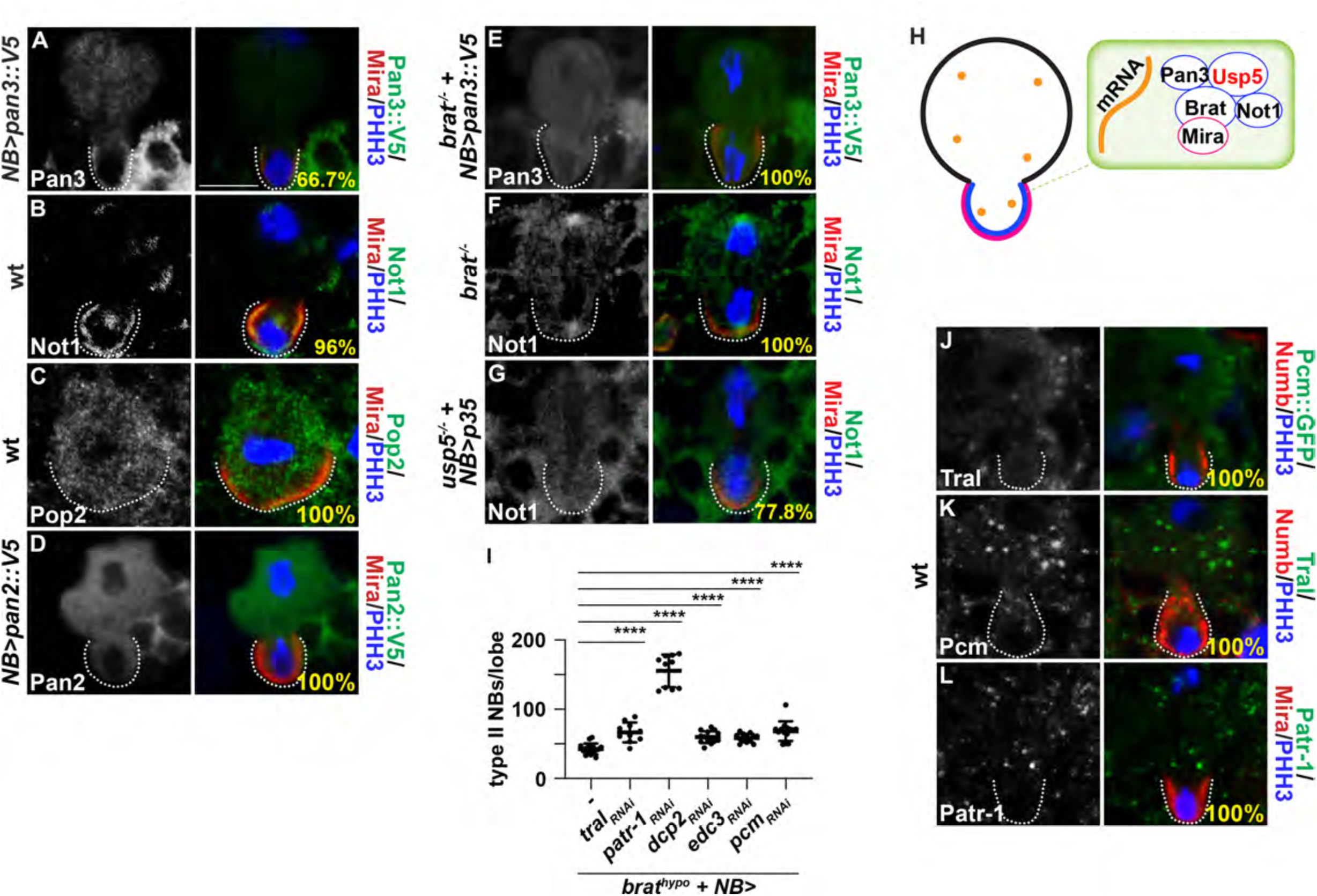
Usp5 promotes assembly of the Brat mRNA decay pre-complex in mitotic neuroblasts. **A-G,** wild-type, *brat*^-/-^ or *usp5^-/-^* larval brains were stained with specific antibodies against the indicated proteins. **H,** Model for assembly of the Brat pre-complex in mitotic neuroblasts. **I,** Quantification of total Type II neuroblasts per lobe of the indicated genotypes. n = 10-16. “-“ indicates no transgene. **J-L,** wild-type larval brains were stained with antibodies against the indicated proteins. Scale bars, 5μm. White dotted lines mark the basal cortex of mitotic neuroblasts. wt: wild-type, *brat^-/-^*: *brat^11^/Df(2L)Exel8040, brat^hypo^*: *brat^DG19310^/^11^, usp5^leon1/leon1^*. Graphs <u>show</u> mean number <u>±</u> standard deviation. P-values: ****<0.0001.

Removal of the 5’ cap structure follows deadenylation ^14,51,52^. Me31B plays an important role in decapping and reduction of *me31B* function is known to enhance the supernumerary Type II neuroblast phenotype in *brat^hypo^* brains ^14,51,52^. This result supports a model that decapping mechanisms are required for Brat-mediated mRNA decay in immature INPs. Consistent with this model, knocking down the function of several genes that encode components of the decapping machinery also enhanced the supernumerary neuroblast phenotype in *brat^hypo^* brains (Fig. 5I). We tested whether these decapping proteins might reside in the Brat-Usp5 pre-complex by examining their localization patterns and functions during asymmetric neuroblast division. We found that decapping proteins are uniformly distributed in the cytoplasm of metaphase and telophase neuroblasts (Fig. 5J-L; Fig. S4A-B). Furthermore, knocking down of the function of these genes alone had no effect on INP commitment in immature INPs (Fig. S4C). These data suggest that decapping proteins are not components of the Brat-Usp5 pre-complex and may instead function downstream of Brat and Usp5 in immature INPs. We conclude Usp5 promotes assembly of the deadenylase-inactive Brat pre-complex in mitotic neuroblasts.

### Mira-binding regulates cell type-specific activation of the Brat-Usp5 complex

A key unanswered question about control of Brat-Usp5-mediated mRNA decay during the transition from a neuroblast state to the onset of INP commitment in newborn immature INPs is the mechanism that activates its ability to induce transcript degradation. Arginine 875, Phenylalanine 916, Asparagine 933 and Asparagine 976 on the top surface of the NHL domain of Brat are required for binding to target mRNAs while Glycine 774 and Tyrosine 829 are essential for binding Mira ^18,31^. Electrophoretic mobility shift assay indicated that mutating Glycine 774 and Tyrosine 829 has little effect on RNA-binding, but Mira overexpression is sufficient to inhibit Brat-mediated repression of reporter expression in S2 cells ^31^. Thus, Mira-binding might regulate activation of the Brat-Usp5 pre-complex in addition to control its localization during asymmetric neuroblast division.

We first assessed whether Mira-binding regulates the RNA-binding activity of the Brat-Usp5 pre-complex by examining the localization pattern of Brat target mRNAs (*dpn* and *klu*) and control mRNAs (*CycE*) relative to the Mira crescent in the basal cortex of mitotic neuroblasts. We observed that *dpn*, *klu* and *CycE* mRNAs predominantly localize in cytoplasmic punctae instead of enriching in the Mira crescent in wild-type metaphase neuroblasts (Fig. 6A; Fig. S5A). Similarly, these transcripts also appeared to localize to punctae throughout the cytoplasm of *brat*-mutant metaphase neuroblasts (Fig. 6B; Fig. S5B). We quantified the frequency of Brat target mRNAs co-localization with Mira in metaphase neuroblasts in wild-type or *brat*-mutant brains by determining the ratio of *dpn* or *klu* mRNA foci to *CycE* mRNA foci in the Mira crescent. This basal mRNA enrichment score indicated that the frequency of *dpn* and *klu* mRNAs co-localizing with Mira was statistically indistinguishable between wild-type or *brat*-mutant metaphase neuroblasts (Fig. 6C). These results show that Brat and its target transcripts do not co-localize at the basal cortex of mitotic neuroblasts and support a model that Brat-Mira binding inhibits Brat-mRNA interactions.

**Figure 6.**
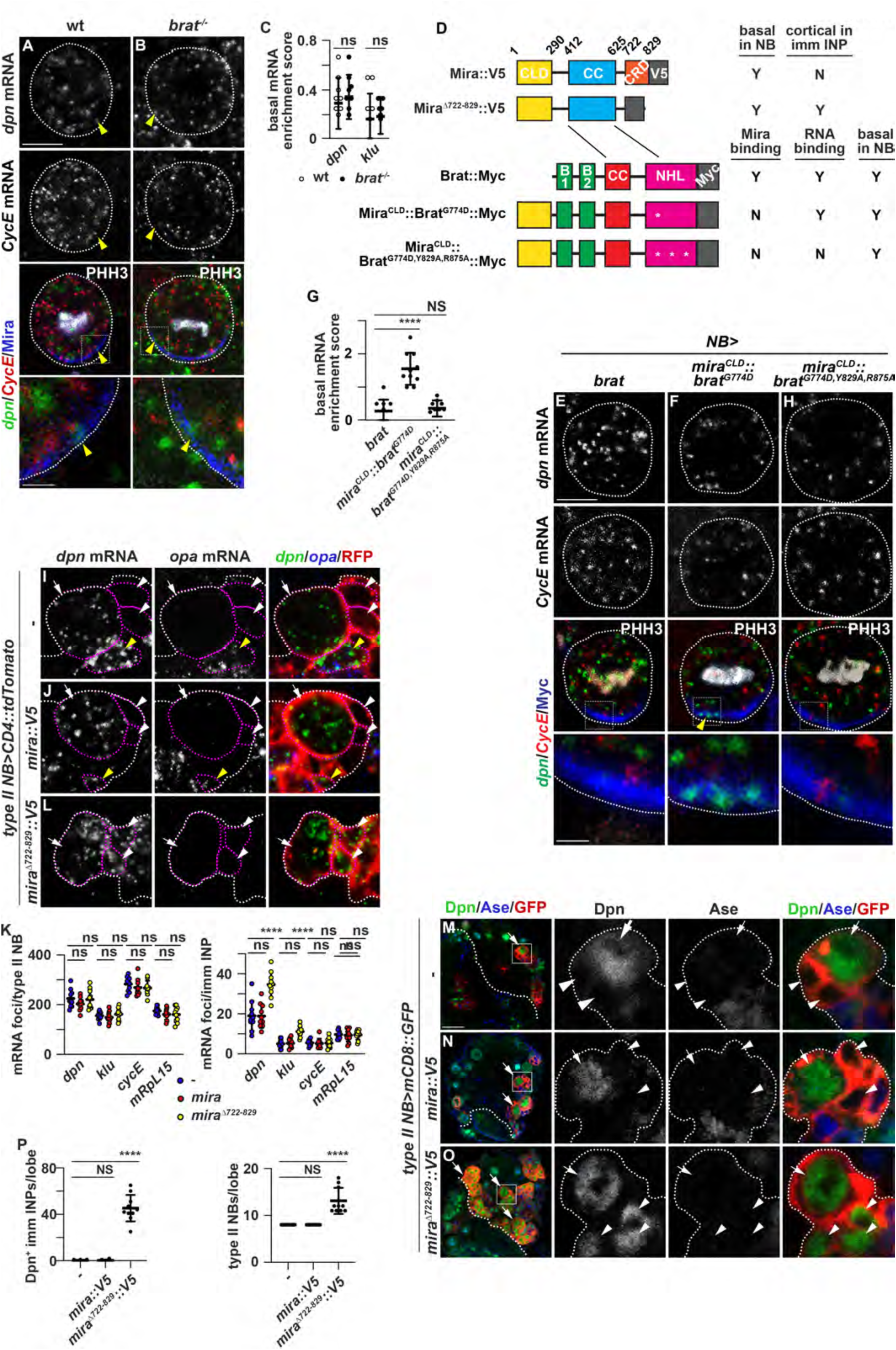
Mira regulates activation of the Brat mRNA decay complex during neuroblast asymmetric division. **A-C,** wild-type or *brat*^-/-^ larval brains were co-hybridized with smFISH probes for *dpn* and *CycE* and counterstained with Mira and PHH3. (C) Quantification of the enrichment of Brat target mRNAs in the basal cortex of mitotic neuroblasts of the indicated genotypes. **D,** The domain structure and experimental results for Mira and Brat encoded by transgenes. B1: B-box 1, B2: B-box 2 CC: Coiled-coil, CLD: Cortical localization domain, CRD: Cargo release domain. Y: Yes. N: No. Domains that mediate Mira and Brat interaction are indicated by the lines. **E-H,** Larval brains overexpressing the indicated transgenes in neuroblasts were co-hybridized with smFISH probes for *dpn* and *CycE* and counterstained with Myc and PHH3. (G) Quantification of the enrichment of *dpn* mRNAs in the basal cortex of mitotic neuroblasts of the indicated genotypes. **I-L,** Larval brains overexpressing the indicated transgenes in Type II neuroblasts were co-hybridized with smFISH probes for *dpn* and *opa* and counterstained with RFP. (K) Quantification of total mRNA foci in Type II neuroblasts or newborn immature INPs in larval brains overexpressing the indicated transgene in the Type II neuroblast lineage. **M-P,** Larval brains overexpressing the indicated transgenes in Type II neuroblasts were stained with antibodies against Dpn, Ase and GFP. (P) Quantification of total Dpn-expressing immature INPs or Type II neuroblasts per brain lobe of the indicated genotypes. Scale bars, 5μm in all images with the following exception. Scale bars, 1μm in enlarged images shown in A-B, E-G. Scale bars, 20μm in low-magnification images shown in M-O. *NB>*: *Wor-Gal4*. *Type II NB>*: *Wor-Gal4,Ase-Gal80*. White dotted lines outline Type II neuroblasts (I-K, M’-O’) or separate the optic lobe from the central brain region (M-O). Magenta dotted lines outline the boundary of cells. White arrows: Type II neuroblast. White arrowheads: newborn immature INP. Graphs show mean ± standard deviation. P-values: ****<0.0001. “ns” indicates no significance.

If Mira-binding sequesters Brat in a non-RNA-binding conformation in mitotic neuroblasts, then mutating amino acid residues in Brat that mediate Mira-binding should bypass this inhibition and increase Brat-mRNA interactions. To increase the specificity of detecting co-localization of Brat and Brat target mRNAs, we targeted Mira-binding-deficient Brat (Brat^G774D^) to the basal crescent in metaphase neuroblasts by fusing the cortical localization domain of Mira (Mira^CLD^) to Brat^G774D^ to generate a Mira^CLD^::Brat^G774D^ chimera (Fig. 6D). We confirmed that full-length Brat and Mira^CLD^::Brat^G774D^ co-localized with Mira in metaphase neuroblasts whereas Brat^G774D^ was present throughout the cytoplasm (Fig. S5C-E). *dpn* and *klu* mRNAs remained in cytoplasmic punctae in metaphase neuroblasts overexpressing full-length Brat, but become enriched in the basal cortex of metaphase neuroblasts overexpressing Mira^CLD^::Brat^G774D^ (Fig. 6E-G; Fig. S5H-J). Importantly, metaphase neuroblasts overexpressing Mira^CLD^::Brat^G774D,Y829A,R875A^ carrying amino acid substitutions at residues that are essential for Brat binding to mRNAs did not show enrichment of *dpn* and *klu* mRNAs in the basal crescent (Fig. 6G,H; Fig. S5J,K). These data strongly suggest that Mira-binding limits the ability of Brat to interact with its target mRNAs in mitotic neuroblasts.

Previous studies have suggested that Mira is degraded in neuroblast progeny destined to differentiate following the dissociation from the cortex and allowing for the release of its cargo proteins ^35,36^. These findings suggest that Mira degradation likely provides a mechanism for cell type-specific activation of the Brat-Usp5 complex in immature INPs. We overexpressed a *UAS* transgene that encodes Mira^Δ722-829^::V5 lacking the cortical dissociation domain (Fig. 6D). Mira^Δ722-829^::V5 is predicted to constitutively associate with the cell cortex of newborn immature INPs and sequester endogenously expressed Brat. We confirmed that overexpressed Mira::V5 and Mira^Δ722-829^::V5 asymmetrically localized in a basal crescent in metaphase neuroblasts and exclusively segregated into the cortex of newborn immature INPs (Fig. S5L,M). Mira::V5 overexpressed in mitotic Type II neuroblasts became displaced into the cytoplasm of immature INPs and had no effect on the decay of either Brat target transcripts (*dpn* and *klu*) and non-Brat target transcripts (*cycE* and *mRpL15*) (Fig. 6I-K; Fig. S5N,O). By contrast, Mira^Δ722-829^::V5 overexpressed in mitotic Type II neuroblasts remained associated with the cortex of immature INPs and resulted in ectopic persistence of Brat target mRNAs but not control non-Brat target mRNAs (Fig. 6K,L; Fig. S5P). Importantly, ectopic persistence of Brat target mRNAs led to ectopic protein expression in immature INPs, driving their reversion into supernumerary Type II neuroblasts (Fig. 6M-P). Together, these data indicate that Mira degradation in immature INPs is required for the activation of the Brat-Usp5 complex for timely onset of INP commitment.

## Discussion

Cell-state transitions must be completed in a timely manner to ensure normal development ^2,5,6^. Rapid and robust clearance of mRNAs is a critical component of the multilayered gene regulatory mechanism that drives timely transitions. RBPs and their downstream effectors have been implicated in many cell-state transitions, but less is known about spatiotemporal control of the assembly and activation RBP-effector complexes. Brat triggers the decay of hundreds of maternally deposited transcripts in the early embryo and neuroblast mRNAs in newborn immature INPs ^14,29,30^. In embryos, we have identified Usp5 as a Brat-interacting protein in embryos that is required for the degradation of Brat target mRNAs. We went on to show that Usp5 promotes the assembly of a Brat-Usp5 pre-complex that lacks deadenylase activity in mitotic neuroblasts (Fig. 7). In parallel, Mira binding to Brat sequesters the Brat-Usp5 pre-complex in an RNA-binding deficient state during asymmetric neuroblast division (Fig. 7). Degradation of Mira in immature INPs then alleviates the inhibition of Brat RNA-binding activity and activates the deadenylase activity of the Brat-Usp5 complex (Fig. 7). We propose that assembly of the pre-complex poises the post-transcriptional regulatory machinery for rapid activation to clear transcripts, thus allowing for timely cell-state transitions.

**Figure 7.**
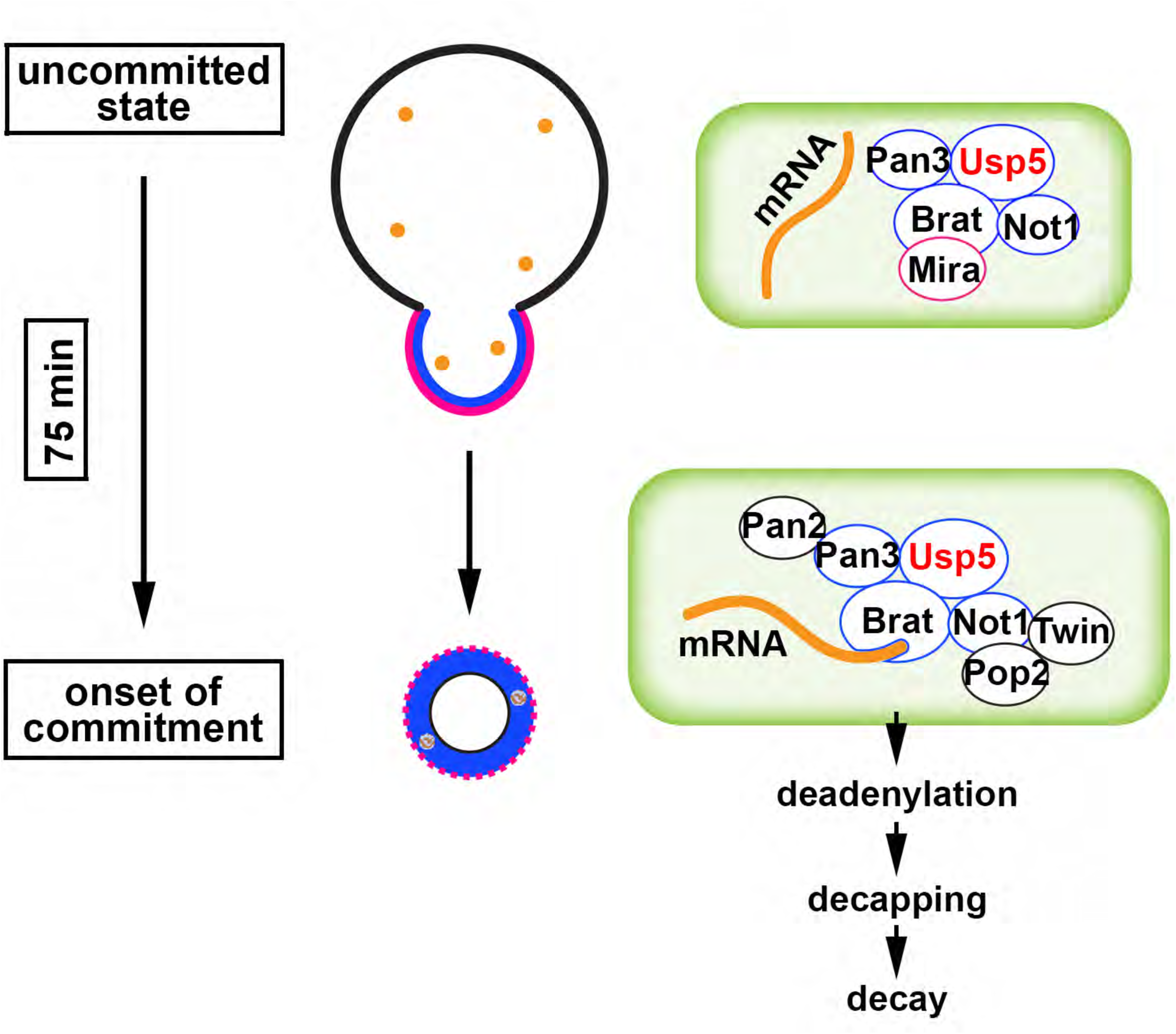
Model for regulation and function of the Brat mRNA decay complex during Type II neuroblast asymmetric division.

### Pre-complex assembly allows for precise spatiotemporal control of mRNA decay machineries

Although the mechanisms by which RNA-binding proteins recognize target transcripts are well-defined, how biologically active mRNA decay complexes are assembled and activated to drive timely mRNA clearance during cell-state transitions is not well-understood ^1-4^. Here we have exploited the known subcellular localization patterns and cell-type-specific function of Brat in the Type II neuroblast lineage to validate Brat-Usp5 complex formation and to define spatiotemporal control of its activity (Fig. 2B-G). Our data suggest that Usp5 functions through the conserved C341 residue to promote the assembly of the Brat-deadenylase pre-complex. Usp5^C341A^ fails to undergo Brat-mediated asymmetric localization and segregation in mitotic neuroblasts (Fig. 2H-K). Overexpressed Usp5^C341A^ cannot substitute for the function of wild-type Usp5 to promote the clearance of Brat target mRNAs in newborn immature INPs (Fig. 3H-K). Furthermore, the scaffolding subunits of RNA deadenylase complexes depend on Brat and Usp5 for asymmetric localization and segregation in mitotic neuroblast (Fig. 5A-H). By contrast, the enzymatic subunits of RNA deadenylase complexes uniformly segregate into both neuroblast progeny. These data led us to propose that Brat and Usp5 form a pre-complex in neuroblasts and addition of the enzymatic subunits to the Brat-Usp5 pre-complex drives rapid activation of deadenylase activity in newborn immature INPs.

We have also shown that Mira is an important contributor to the spatiotemporal control of Brat-mediated mRNA decay. Brat target mRNAs co-localize with transgenic Brat protein tethered basally by the cortical localization domain of Mira, but not with transgenic Brat protein sequestered there by Mira, strongly suggest that Mira-binding limits the ability of Brat to bind its target mRNAs (Fig. 6B-F; Fig. S6H-K). Our observations are consistent with previously reported findings that Mira overexpression suppresses Brat-3’UTR reporter activity in S2 cells ^29^. Because Brat target mRNAs include transcripts that encode factors required to maintain Type II neuroblasts in an undifferentiated state, these results imply that Mira promotes neuroblast maintenance by limiting Brat-mediated mRNA decay in mitotic neuroblasts. Mira becomes displaced from the cortex to the cytoplasm and degraded in neuroblast progeny destined to differentiate. Cortical retention of Mira prevents release of the Brat pre-complex into the cytoplasm of neuroblast progeny leading to their reversion to supernumerary neuroblasts (Fig. 6M-P). Thus, sequestration of the Brat pre-complex in the cortex of mitotic neuroblasts by Mira might inhibit assembly of the full Brat mRNA decay complex. We previously demonstrated that the RNA-binding protein Tis11 likely contributes to Brat recognition of high-affinity target transcripts and the overall robustness of Brat-mediated mRNA decay ^14^. Tis11 is expressed at a very low level endogenously, and overexpressed transgenic Tis11 is exclusively detected in the cytoplasm of newborn immature INPs. Pop2 and Pan2 are enzymatic subunits of the CCR4-Not1 and Pan2-Pan3 deadenylase complexes ^52,53^. Together, these data suggest that biologically active Brat-Tis11-Usp5 mRNA decay complex exists in the cytoplasm of newborn immature INPs and that Mira-mediated sequestration of Brat-Usp5 pre-complex likely exerts an inhibitory effect on assembly and activation the Brat complex.

### A conserved role for Usp5 as a scaffolding protein

Hydrolysis of ubiquitin from ubiquitylated proteins or unanchored polyubiquitin chains into free monoubiquitin is the most well-recognized mechanism by which Usp5 regulates biological processes ^41,44,45,54^. The ubiquitin-specific processing protease domain of Usp5 exerts deubiquitinase activity, but other domains directly or indirectly influence its catalytic activity. Structural analysis of full-length Usp5 led to discovery of the cryptic nZn-UBP that tightly interacts with the enzymatic domain and is indispensable for its catalytic activity despite lacking enzymatic activity ^46^. UBAs in the C-terminus indirectly contribute to deubiquitinase activity of Usp5 by binding ubiquitin ^55,56^. Studies of the vertebrate immune response suggest that Usp5 can function as a scaffold to promote the assembly of multi-subunit protein complexes independent of its enzymatic activity. Usp5 regulates the Type I interferon (IFN) response against invading RNA and DNA viruses by promoting interactions between the E3 ubiquitin ligase complex and ubiquitin via UBAs ^57^. Usp5 functions through the ubiquitin-specific processing protease domain to recruit the E3 ubiquitin ligase complex to promote polyubiquitination and autophagic degradation components of certain inflammasome components in vertebrates ^58^. However, the scaffolding role of Usp5 in promoting protein complex assembly and activity remains poorly studied.

Our data suggest that Usp5 functions as a scaffolding protein independent of its enzymatic activity to promote Brat mRNA decay pre-complex assembly during asymmetric neuroblast division. Usp5 functions through the conserved C341 residue in the ubiquitin-specific processing protease domain to undergo Brat-dependent asymmetric segregation into newborn immature INPs and to promote INP commitment (Fig. 2I-K; Fig.3E-G). Identical to the *usp5*-null (*usp5^leon1^*^/*Df*^) allelic combination, the *usp5^hypo^*(*usp5^leon1^*^/*leon2*^) also reduced INP commitment in *brat^hypo^* brains leading to enhancement of the supernumerary neuroblast phenotype (Fig. 3J, L). Both the *usp5^leon1^*and *usp5^leon2^* alleles have a single amino acid substitution in the cZn-UBP domain (Fig. 3A), and deletion of this domain does not significantly affect the rate of Usp5 hydrolysis of ubiquitin ^42,46^. Structural analyses suggest that the cZn-UBP domain contains a deep binding pocket while the linkers on either side are disordered and lengthy, allowing for considerable flexibility to accommodate substrates with a variety of structural ensembles within the vicinity of the ubiquitin-specific processing protease domain ^46,47^. Thus, we propose that Usp5 functions through the cZn-UBP domain and ubiquitin-specific processing protease domain to promote the assembly of the Brat pre-complex binding to Brat. The C-terminal UBA interacts extensively with the ubiquitin-specific processing protease domain, and its loosely tethered nature allows for considerable conformational mobility relative to the ubiquitin-specific processing protease domain, allowing for binding to a variety of proteins. In the future, it will be interesting to test whether the C-terminal UBA of Usp5 also plays a role in bridging Brat and scaffolding components of deadenylase complexes.

### Usp5 regulates gene expression at multiple levels during development and homeostasis

Its deubiquitinase activity appears to play a key role in driving most Usp5-regulated biological processes. Emerging evidence suggests that Usp5 might regulate gene expression at both the protein and mRNA levels. Usp5 suppresses autophagy in fly and vertebrate cells by maintaining the autophagy initiator protein Atg1 at a low level ^59^. The C-terminal UABs are required for Usp5 binding to Atg1 but the conserved Cysteine in the ubiquitin-specific processing protease domain is dispensable for Usp5-Atg1 interactions. Loss of *usp5* function leads to increased levels of ubiquitinated Atg1 protein, but there is no evidence indicating that Usp5 directly modulates the ubiquitination of Atg1. Notably, loss of *usp5* function also dramatically increased levels of *atg1* mRNAs suggesting that Usp5 might regulate *atg1* transcript stability. In this study, we have demonstrated that loss of *usp5* function increases Notch target gene expression at the protein and mRNA levels (Fig. 4H-J, N-P). Our data further indicate that Usp5 is a key scaffold that facilitates assembly of the Brat mRNA decay complex that targets Notch downstream-effector gene transcripts for decay. We speculate that Usp5 might suppress autophagy by maintaining low levels of Atg1 protein via the decay of *atg1* mRNAs. We propose that Usp5 likely functions as a multi-functional regulator of gene expression by controlling proper protein and mRNA levels.

## Acknowledgements

We would like to thank Alexa Gonzalez for technical assistance. We would like to thank R. Warton, J. Skeath, E. Whale, C.T. Chien, G. Rogers, S. Barbee, N. Sokol and A. Nakamura for reagents and the Bloomington *Drosophila* Stock Center and the Vienna *Drosophila* Resource Center for fly stocks. We would like to thank Genetivision for generating transgenic fly lines. We would like to thank members of the Lee lab for comment and feedback. This work was supported by grants from the Canadian Institute for Health (PJT-15970 and PJT-19012 to H.D.L) and the National Institute of Neurological Disorders and Stroke grants (R01NS111647 and R01NS107496 to C.-Y.L).

## Methods

### KEY RESOURCES TABLE

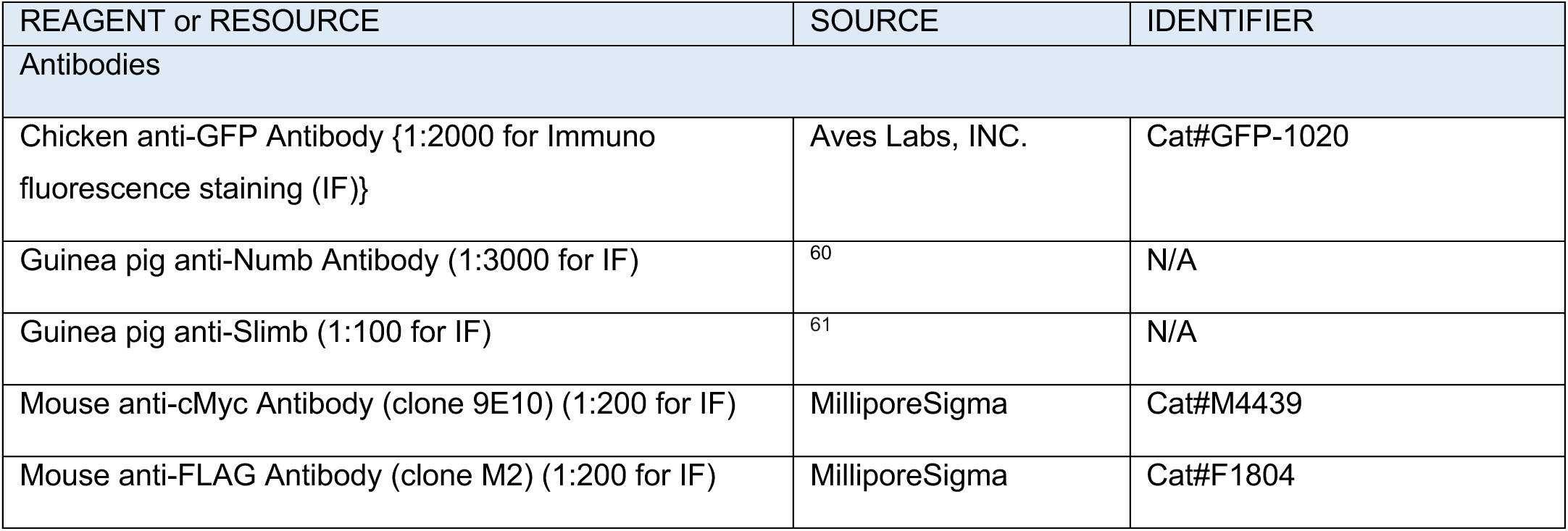
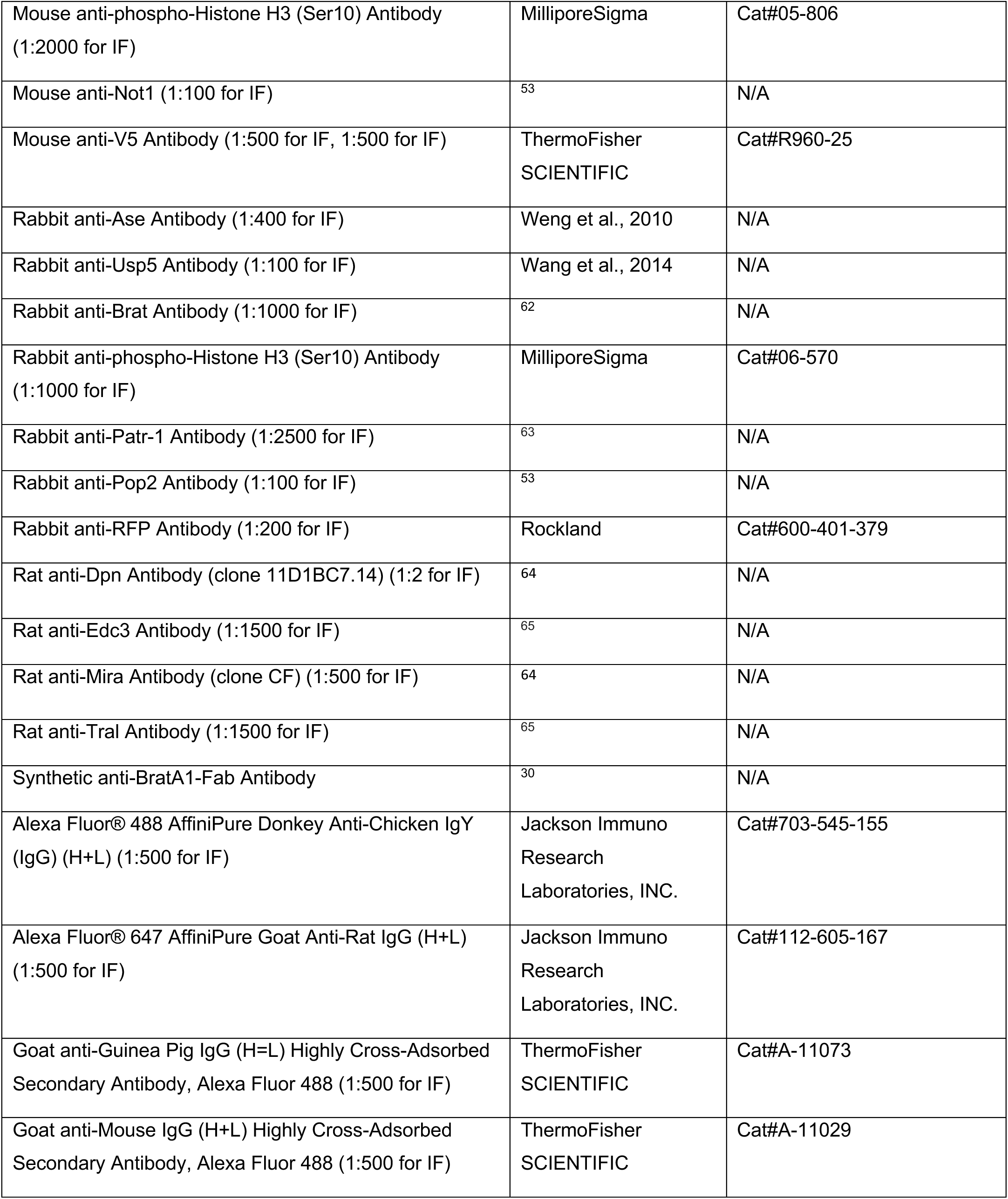
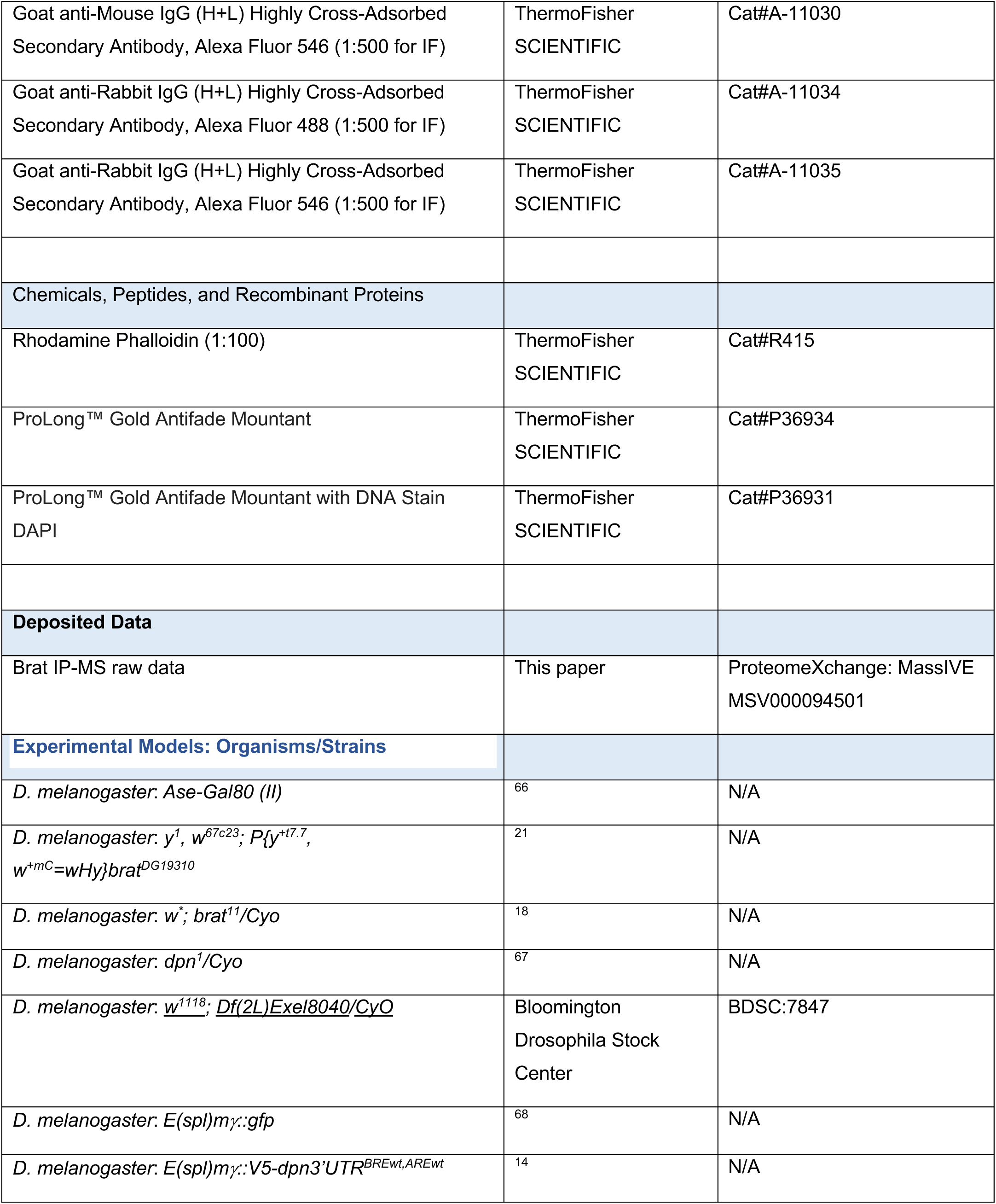
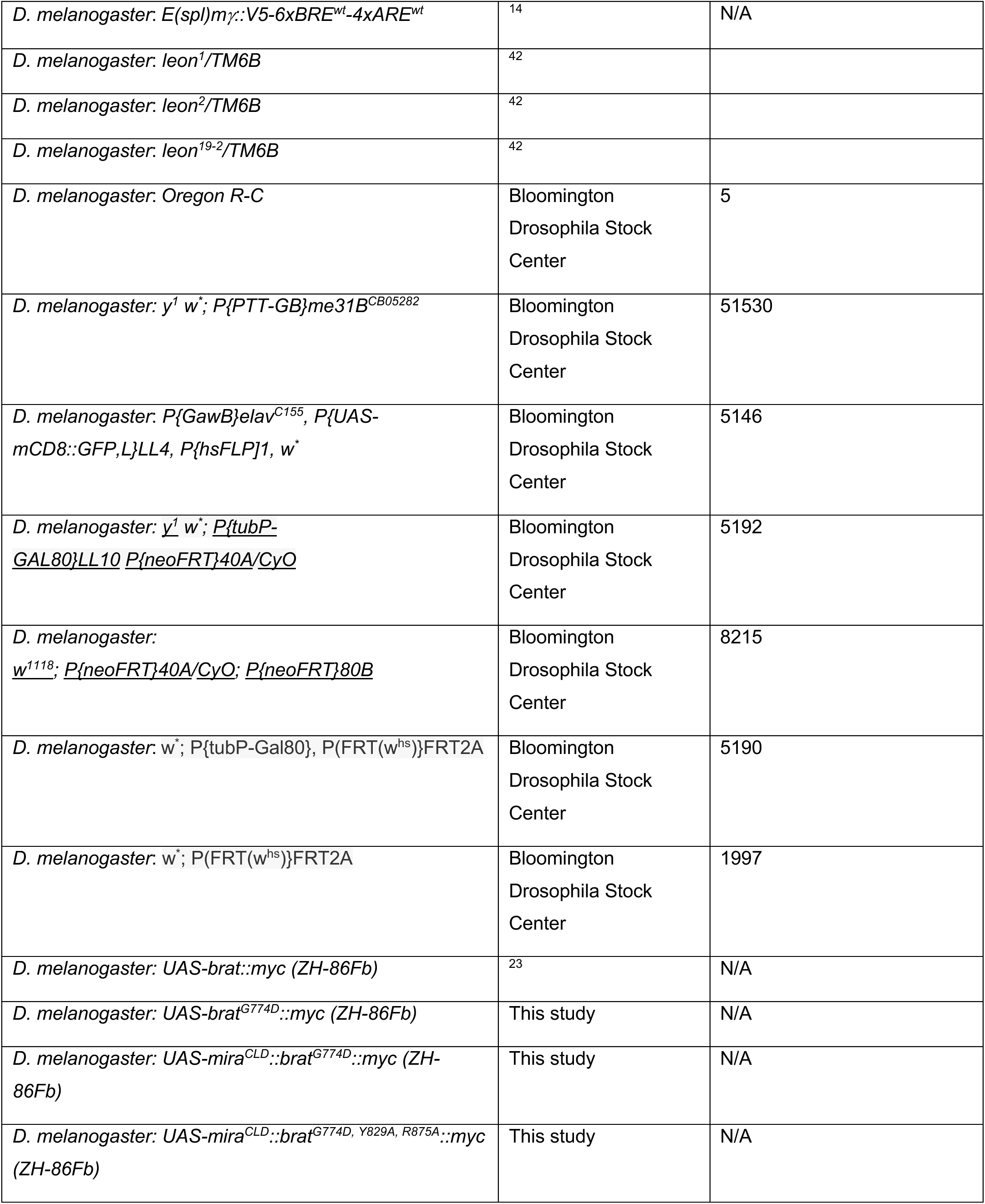
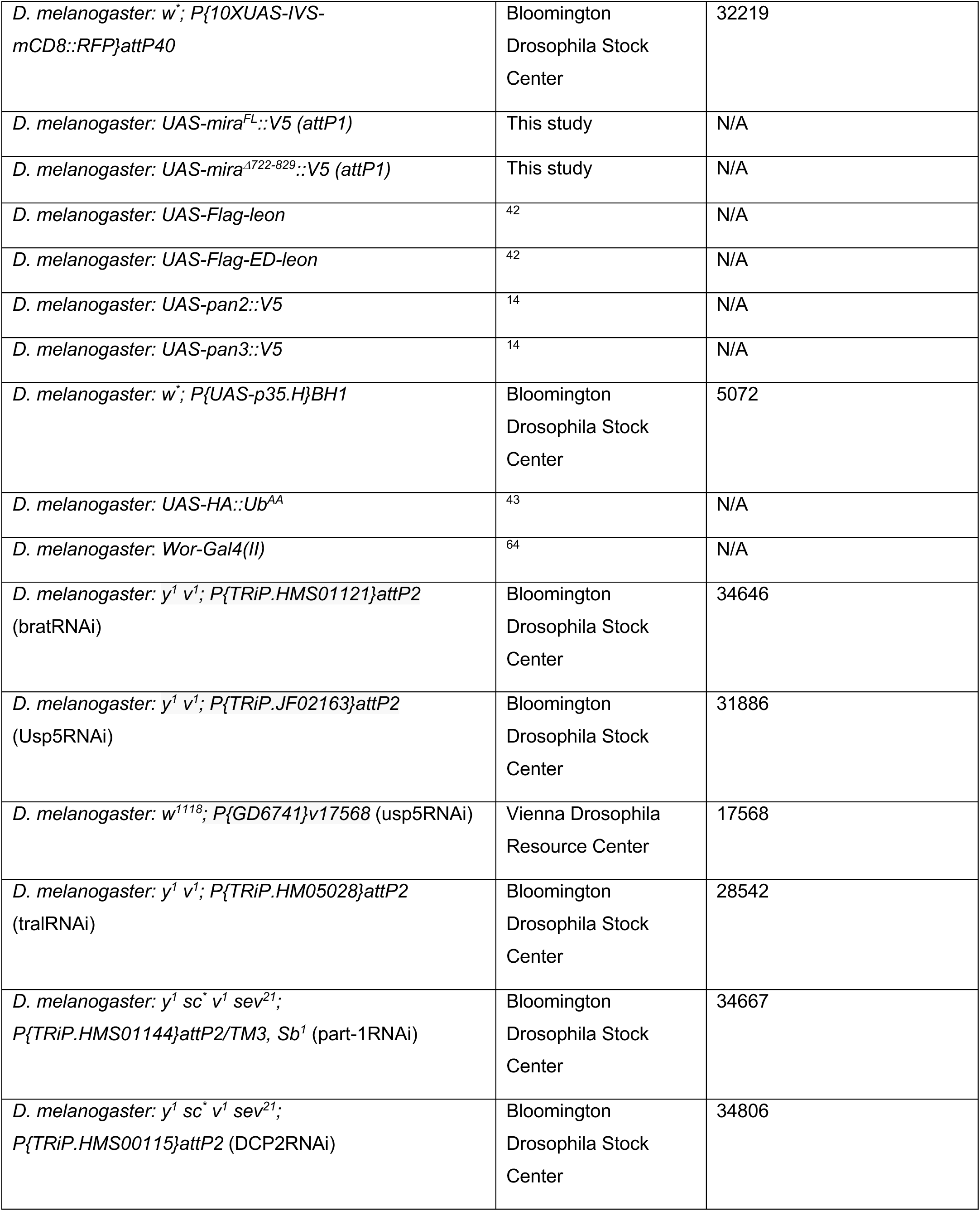
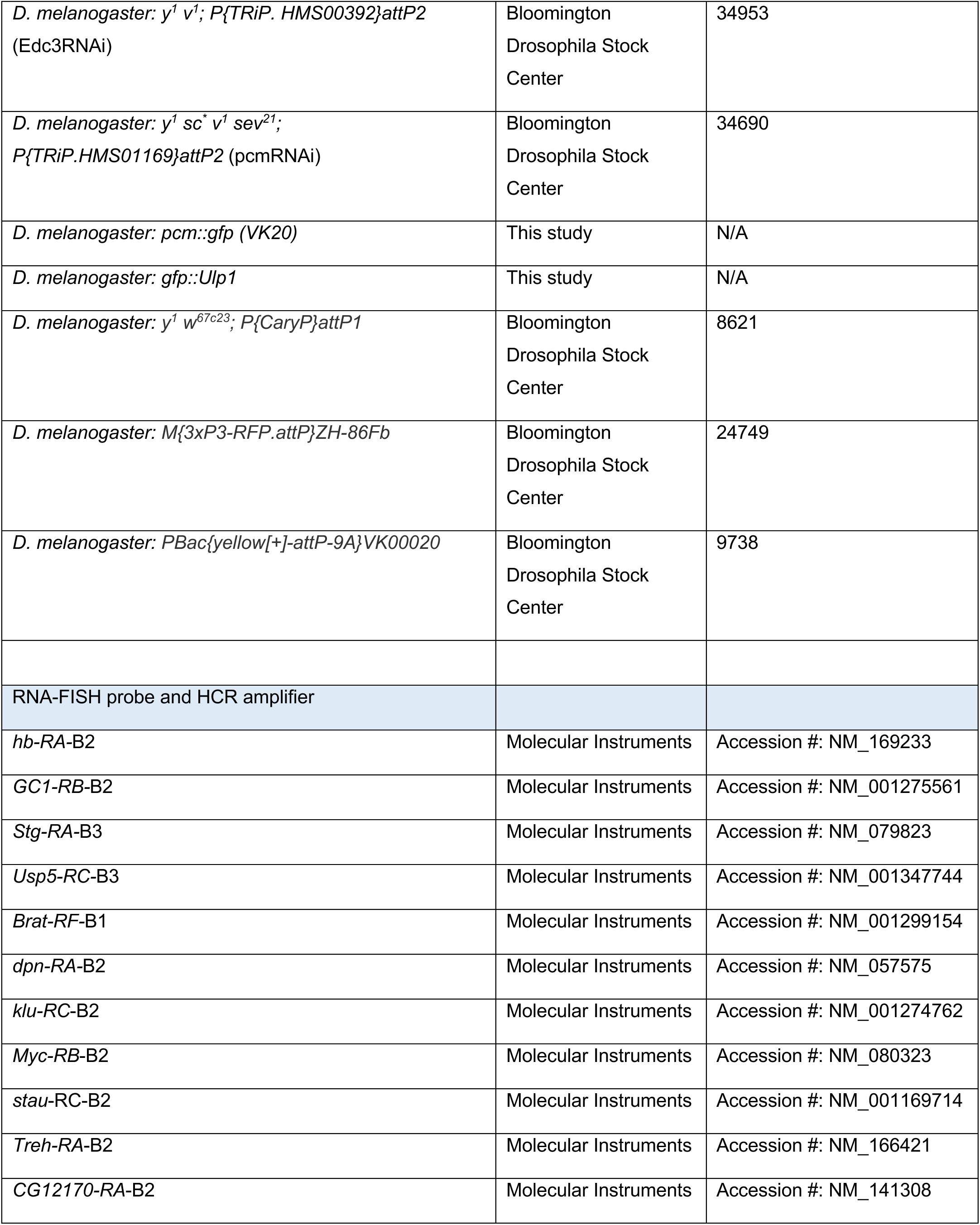
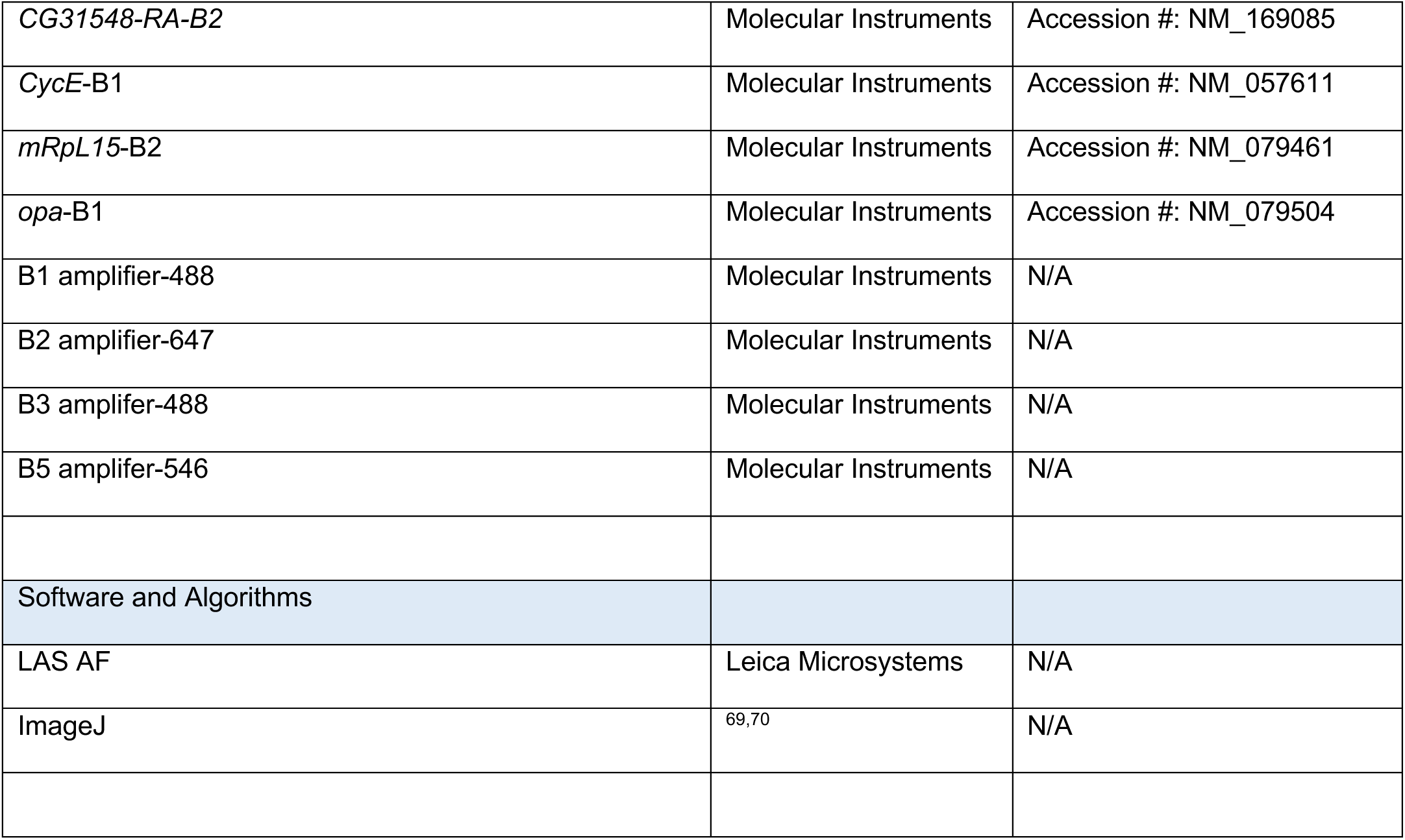

#### Fly culture condition

All fly stocks were maintained on standard *Drosophila* culture media at 25°C. Eggs were collected at 25°C for 3 hours period for obtaining 0-3 hours embryo extracts, a 2-hour period for mRNA staining, or an 8-hour period for larval brain experiments. Larvae were cultured on caps containing *Drosophila* culture media at 25°C or 33°C. Larvae were cultured at 25°C for 96 hours after larval hatching (ALH) for experiments of Fig. 2B-G, Fig. S4A-B, Fig. 5B-C, J-L, Fig. S5A-B, Fig. 6A-C. To drive high levels of transgene expression in late third instar larvae, larvae were cultured at 33°C for 72 hours ALH (experiments of Fig. 2H-K, Fig. S2A-C, Fig. 3B-L, Fig. S3O-Q, Fig. 4A-B, K-M, Q-S, Fig. S4C, Fig. 5A, D-G, I, Fig. S5N-P, Fig. 6I-P). To limit transgenes in neuroblast and surrounding progenies, larvae were cultured at 25°C until 64 hours ALH and shift temperature to 33°C for the last 20 hours before dissection (Fig. S5C-M, Fig. 6E-H).

For MARCM analysis (Fig. S3B-N, Fig. 4C-J, N-P) embryos or larvae were incubated at 37°C for 90 min to induce clones and then, the heat-shocked animals were cultured at 25°C prior to dissection.

#### Fly genetics

The *brat^G774D^*, *mira^FL^*, *mira^Δ722-829^*, and *mira::brat* chimeric transgenes were cloned into the *p{UAST}attB* vector. The *pcm::gfp(g)* clone was generated by inserting GFP sequence in frame to *pcm* in the BAC clone (CH322-140C01). The transgenic fly lines were generated via ϕC31 integrase-mediated transgenesis by using *y^1^w^67c23^; P{CaryP}attP1* (for *UAS-mira* constructs), *M{3xP3-RFP.attP}ZH-86Fb* (for *UAS-brat* constructs), and *PBac{yellow[+]-attP-9A}VK00020* (for *pcm::gfp*).

#### Immunoprecipitation and mass spectrometry

Methods were as described in Laver et al ^30^. 0-3-hour old embryos were collected, dechorionated and crushed in a minimal volume of lysis buffer (150 mM KCl, 20 mM HEPES-KOH pH 7.4, 1 mM MgCl2, 0.1% Triton X-100, supplemented with protease inhibitors and 1 mM DTT), cleared by centrifugation for 15 min at 4°C and 20,000 g, and stored at -80°C. Immediately prior to immunoprecipitation (IP), sample was diluted to 15 mg/mL protein concentration with lysis buffer. For IP, 800 μL of diluted lysate was mixed with or without 350 μg/μL RNase A, 40 μL of anti-Flag M2 beads (Sigma) that were pre-loaded with 20 μg of either anti-BratA1 Fab antibody or control C1 Fab antibody. IPs were incubated for 3-hour at 4°C with end-over-end rotation. Beads were washed four times with lysis buffer, twice with lysis buffer lacking Triton X-100, then transferred to new tubes and washed twice more with lysis buffer lacking Triton X-100 to remove all traces of detergent. Bound proteins were eluted by tryptic digest: beads were resuspended in 200 μL of 50 mM ammonium bicarbonate pH 8, supplemented with 2 μg of trypsin, and incubated overnight at room temperature with end-over-end rotation. The following day, the digested supernatant was recovered, and beads were washed once with an additional 200 μL of 50 mM ammonium bicarbonate to collect any residual eluted material. The two supernatants were pooled and dried by speed-vac. Liquid chromatography–tandem mass spectrometry (LC-MS/MS) was performed using a Thermo Q-Exactive HF quadrupole-Orbitrap mass spectrometer (Thermo Scientific) following previously described methods ^71-73^. Three biological replicates of Brat IPs and control IPs were performed, in both the presence and absence of RNase A.

To identify Brat-interacting proteins, we used the ProHits software package ^74^ to perform <u>S</u>ignificance <u>A</u>nalysis of <u>INT</u>eractome (SAINT), comparing RNase-treated Brat IP versus control IP samples, and non-RNase-treated Brat IP versus control IP samples. Specifically, SAINT input files were generated using the ‘‘TPP iProphet’’ search engine option in ProHits, filtering for iProphet probability <u>≥</u> 0.9 and number of unique peptides < 2. SAINTexpress (exp3.3) was run using the ProHits interface, including only detected Drosophila proteins, with the following settings: number of compressed controls = 3; burn-in period, nburn = 2000; iterations, niter = 5000; lowMode = 1; minFold = 1; normalize = 1; nCompressBaits = 2. Identified proteins were defined as RNA-independent or RNA-dependent Brat-interacting proteins, if, in the analyses of the respective samples, they achieved a SAINT score R ≥ 0.9 and a Bayesian false discovery rate (BFDR) < 0.05.

#### Immunofluorescent staining

Larval brains were dissected in PBS and fixed in 100 mM PIPES (pH 6.9), 1 mM EGTA, 0.3% Triton X-100 and 1 mM MgSO4 containing 4% formaldehyde for 23 minutes. Fixed brain samples were washed with PBST containing PBS and 0.3% Triton X-100. After removing fix solution, samples were incubated with primary antibodies for 3 hours at room temperature. 3 hours later, samples were washed with PBST and then incubated with secondary antibodies overnight at 4°C. On the next day, samples were washed with PBST and then equilibrated in ProLong Gold antifade mount (ThermoFisher Scientific).

Brains were mounted with dorsal surface up and ventral surface down. Confocal images were acquired on a Leica SP5 scanning confocal microscope (Leica Microsystems Inc). To quantify the number of Type II neuroblasts, images of a brain lobe were sequentially taken at every 1.5 μm dorsal to ventral.

More than 10 brains per genotype were used to obtain data in each experiment. Antibodies are listed in the Key Resources Table.

#### Hybridization Chain Reaction (HCR) and immunofluorescent staining

mRNA in the embryo, the larval brain, and the adult ovary were quantified by performing in situ HCR v3.087 ^75^. cDNA sequences were obtained from NCBI to generate HCR probes. HCR probes are listed in the Key Resources Table.

For mRNA staining in early-stage embryos, parental males and females were cultured at 25°C for 6 days before egg collection. Eggs were collected at 25°C for 2 hours and then incubated at 25°C for an additional 1.5 hours. Embryos were dechorionated with 3% sodium hypochlorite for 2 minutes and fixed in heptane/1xPBS containing 4 % formaldehyde (1:1) for 20 minutes. Fixed embryos were devitellinized by shaking in methanol. Samples were pre-hybridized with hybridization buffer (10% formamide, 5×SSC, 0.3% Triton X-100 and 10% dextran sulfate) at 37°C for 1 hour, 5 nM HCR probes (Molecular Instruments) were added and embryos were then incubated at 37°C overnight. After hybridization, samples were washed with washing buffer (10% formamide, 5× SSC, 0.3% Triton X-100) and then incubated with amplification buffer (5× SSC, 0.3% Triton X-100 and 10% dextran sulfate) at 25°C for 30 min. Once samples were equilibrated in amplification buffer, samples were mixed with 3 μM of imager hairpins and incubated at 25°C for overnight. The next day, samples were washed with PBST.

Third instar larval brains or adult ovaries (5 days after adult fly eclosion) were dissected in PBS and fixed in 100 mM PIPES (pH 6.9), 1 mM EGTA, 0.3% Triton X-100 and 1 mM MgSO4 containing 4% formaldehyde for 23 minutes. Fixed samples were washed with PBST containing PBS and 0.3% Triton X-100. The HCR reaction was performed as described above. If antibody staining was required, samples were re-fixed in 100 mM PIPES (pH 6.9), 1 mM EGTA, 0.3% Triton X-100 and 1 mM MgSO4 containing 4% formaldehyde for 15 min to initiate immunofluorescent staining procedures. To quantify mRNA foci in Type II neuroblasts and newborn immature INPs, images of cells were sequentially taken at every 0.5 μm. More than 10 brains per genotype were used to obtain data in each experiment.

#### Quantification and Statistical Analysis for IF and HCR

To quantify the number of Type II neuroblasts in a larval brain lobe, Type II neuroblasts were identified by Dpn^+^Ase^-^ expression and > 9 μm cell diameter. Dpn^+^Ase^-^ cells < 8 μm were classified as immature INPs aberrantly expressing Dpn in this quantification. The number of Type II neuroblasts were manually counted using confocal images.

To quantify the number of maternal mRNA foci in an early-stage embryos, the exact developmental stage was identified using the *tll* mRNA expression and localization pattern. HCR signals of maternal mRNAs in a 10μm^2^ region in the posterior half of each embryo were manually counted using confocal images (Figure 1).

To quantify the number of mRNA foci in Type II neuroblasts and in newborn immature INPs, Type II neuroblasts were identified by the presence of Dpn (in Figure 3A and S3E) or absence of *opa* mRNA foci with cell size (cell diameter > 9 μm) in Type II neuroblast clones while newborn immature INPs were identified by the absence of *opa* mRNA foci and cell location (the most proximal cell to the mother Type II neuroblast). Dpn-positive small cells in Figure 3A and S3E were INPs. *opa* mRNA-positive cells in other images were INPs and their progeny in Type II neuroblast clones. HCR signals in individual cells were manually counted using confocal images.

To quantify the *dpn* or *klu* mRNAs localized in the basal cortex of mitotic neuroblasts, HCR signals of *dpn*, *klu* and *cycE* mRNAs overlapping with the Brat::Myc signal were manually counted using confocal images. The number of *dpn* or *klu* mRNA foci were normalized to the number of *CycE* mRNA foci.

Image J software was used to measure the intensity of E(spl)my-V5 reporter signal. DAPI single channel confocal images were used to identify the nucleus of individual cells, and the pixel intensities of V5 and DAPI were assessed using individual channel confocal images of the same optical section. The mean of the V5 signal intensity was normalized to the mean of the DAPI signal intensity.

The number of biological replicates (n) is indicated in each figure legend and the standard deviation is indicated by the error bars. All statistical analysis for two samples were performed using Wilcoxon signed-rank test. All statistical analysis for more than three samples was performed by using Ordinary one-way ANOVA, a p-value < 0.05, <0.01, <0.001 and <0.0001 is indicated by (*), (**), (***), and (****) respectively in figures.

## Figure Legends

**Supplemental Figure 1.**
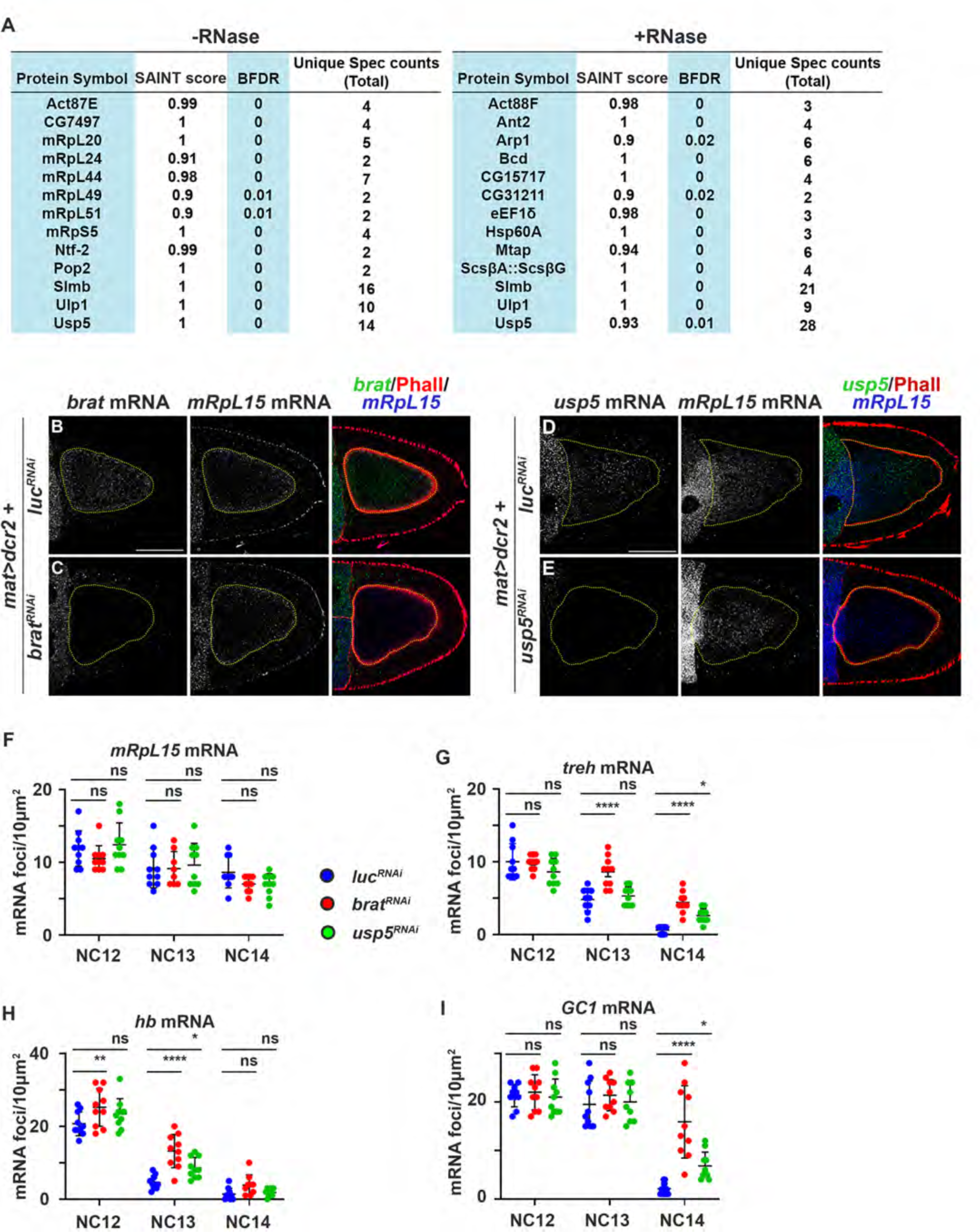
Identification of new Brat interactors in the fly embryo. **A,** A list of highest-confidence Brat interacting proteins identified from 0–3-hour embryo extracts by IP-MS. See Table S1 for complete results. **B-E,** Adult ovary with *brat*, *mRpL15* or *usp5* knocked down were co-hybridized with smFISH probes for *brat, mRpL15* or *usp5* and counterstained with Phalloidin. Scale bars, 50μm. The *P{GD6741}* transgene was used to knock down *usp5* function. **F-I,** Quantification of total mRNA foci in 10 μm^2^ in NC12-14 embryos of indicated genotypes.

**Supplemental Figure 2.**
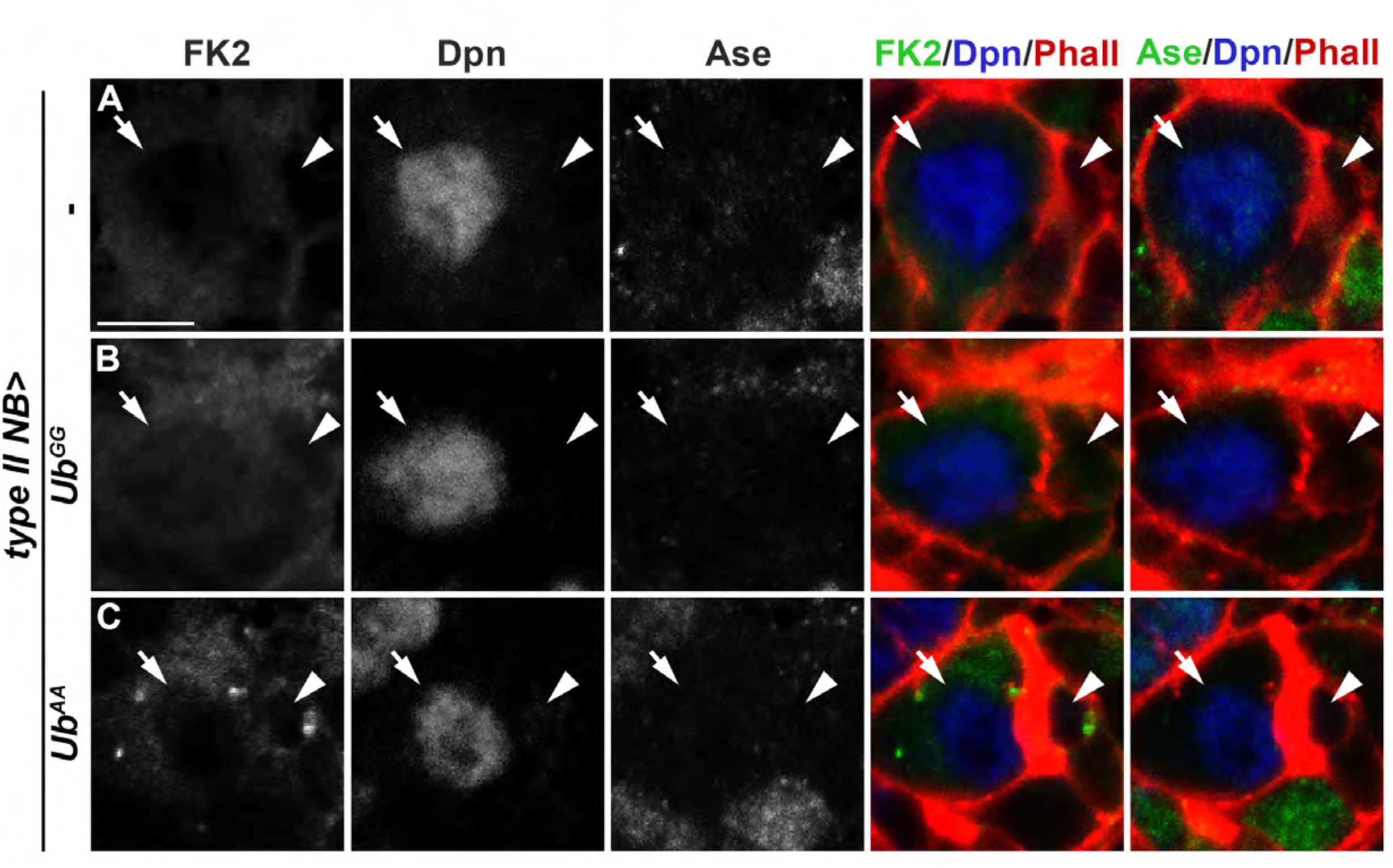
Overexpression of Ub^AA^ leads to ectopic polyubiquitin accumulation. **A-C,** Larval brains overexpressing the indicated transgenes were stained with antibodies against FK2, Dpn, Ase and Phalloidin. The FK2 antibody specifically reacts with polyubiquitin chain. Scale bars, 5μm. White arrows: Type II neuroblast. White arrowheads: newborn immature INP.

**Supplemental Figure 3.**
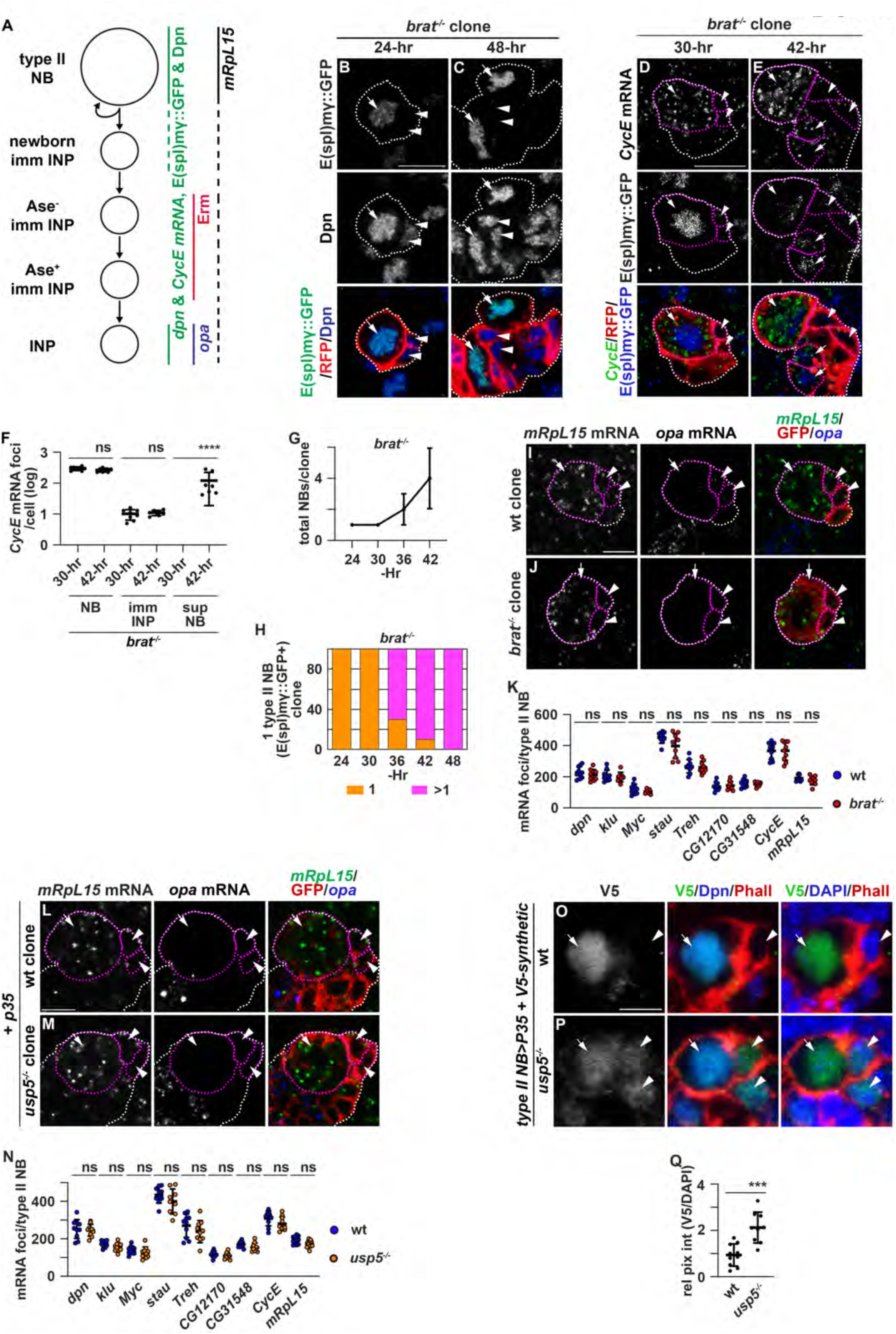
Usp5 promotes clearance of Brat target transcripts in immature INPs. **A,** Schematic of gene expression patterns in the Type II neuroblast lineage. **B-H**, Larval brains carrying an *E(spl)mγ::GFP* transgene and containing RFP-marked *brat^-/-^*mosaic clones were aged for the indicated length of time following induction and stained with the indicated markers. (F) Quantification of *CycE* mRNA foci in indicated cell types in *brat^-/-^* Type II neuroblast clones. n = 10. (G) The total number of neuroblasts *brat^-/-^*mosaic clones. (H) The percentage of *brat^-/-^* Type II neuroblast clones containing >1 neuroblast per clone. **I-K**, Larval brains carrying GFP-marked wild-type or *brat^-/-^* Type II neuroblast mosaic clones aged for 24 hours following induction were co-hybridized with the indicated smFISH probes and were counterstained with an antibody against GFP. (K) Quantification of the indicated mRNA foci per Type II neuroblast of the indicated genotype. **L-N**, Larval brains carrying GFP-marked wild-type or *usp5^-/-^*Type II neuroblast mosaic clones aged for 24 hours following induction were co-hybridized with the indicated smFISH probes and were counterstained with an antibody against GFP. (N) Quantification of the indicated mRNA foci per Type II neuroblast of the indicated genotype. **O-Q,** wild-type or *usp5^-/-^* larval brains carrying a *V5-synthetic* reporter transgene and overexpressing P35 were stained with specific antibodies against V5, Dpn, DAPI and Phalloidin. (Q) Quantification of relative reporter expression in newborn immature INPs of indicated genotypes. The relative reporter expression levels were determined by calculation the ratio of V5 to DAPI. n = 8-10 brains. Scale bars, 5μm. White dotted lines outline Type II neuroblast clones. Magenta dotted lines outline the boundary of cells. White arrows: Type II neuroblast. White arrowheads: newborn immature INPs. Yellow arrows: INP. Graphs show mean ± standard deviation. P-values: ***<0.001, ****<0.0001. “ns” indicates no significance. wt: wild-type, *brat^-/-^*: *brat^11/11^*, *usp5^-/-^*: *usp5^leon1/leon1^*.

**Supplemental Figure 4.**
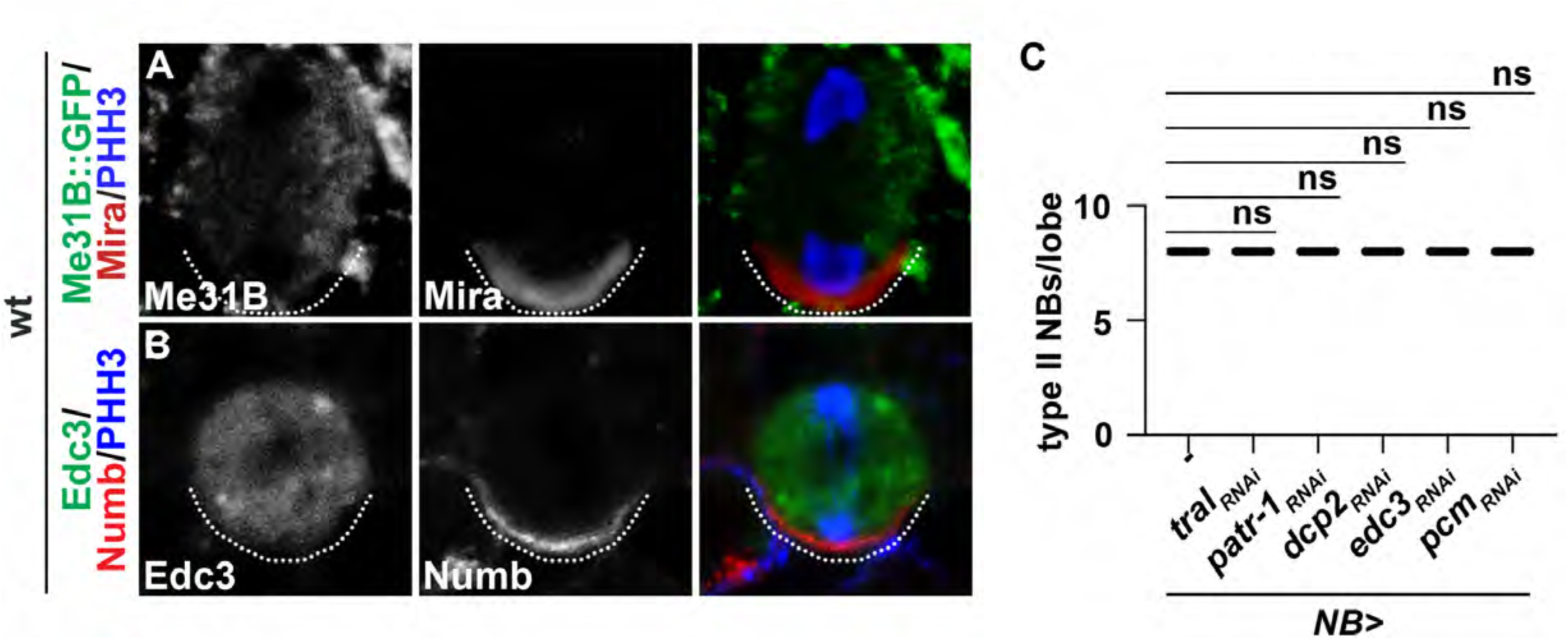
Decapping proteins are uniformly distributed in the cytoplasm of mitotic neuroblasts. **A-B,** wild-type larval brains alone or carrying a *Me31B-GFP* transgene were stained with specific antibodies against the indicated proteins. **C,** Quantification of total Type II neuroblasts per lobe of the indicated genotypes. n = 10. Graphs show mean ± standard deviation. “ns” indicates no significance.

**Supplemental Figure 5:**
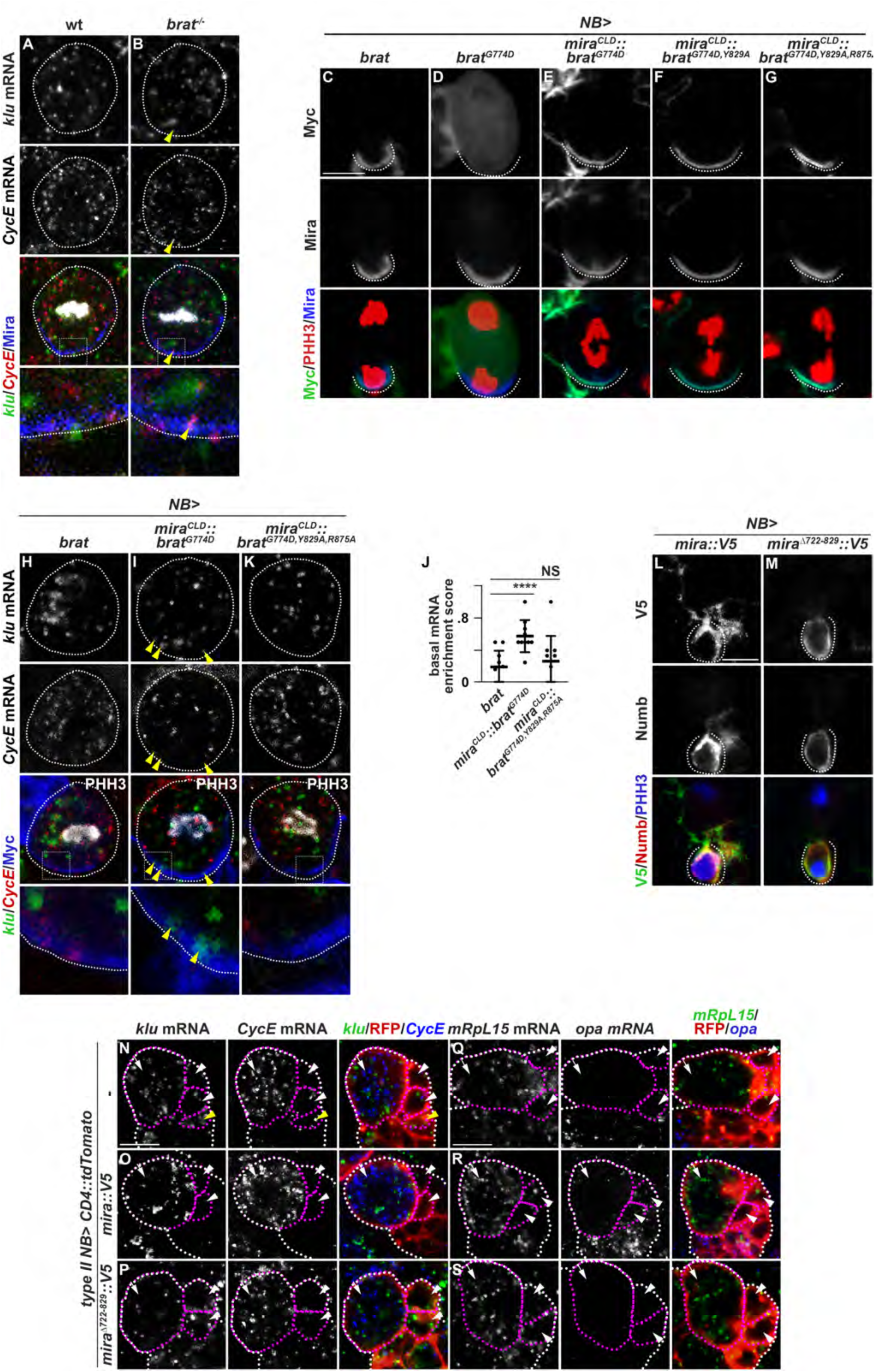
Mira regulates activation of the Brat mRNA decay complex. **A-B,** wild-type or *brat*^-/-^ larval brains were co-hybridized with smFISH probes for *klu* and *CycE* and counterstained for Mira and PHH3. **C-G,** Larval brains overexpressing the indicated transgene in neuroblasts were stained for Myc, Mira and PHH3. **H-K,** Larval brains overexpressing the indicated transgene in neuroblasts were co-hybridized with smFISH probes for *klu* and *CycE* and counterstained for Myc and PHH3. (J) Quantification of the enrichment of *dpn* mRNAs in the basal cortex of mitotic neuroblasts of the indicated genotypes. **L-M,** Larval brains overexpressing the indicated transgene in neuroblasts were stained with antibodies against V5, Numb and PHH3. **N-S,** Larval brains overexpressing the indicated transgenes in Type II neuroblasts were co-hybridized with smFISH probes for *klu*, *CycE*, *opa* and *mRpL15* and counterstained with RFP. Scale bars, 5μm except 1μm in enlarged images shown in the bottom panels of A,B,H,I,K. *NB>*: *Wor-Gal4*. *Type II NB>*: *Wor-Gal4,Ase-Gal80*. White dotted lines outline Type II neuroblast clones. Magenta dotted lines outline the boundary of cells. White arrows: Type II neuroblast. White arrowheads: newborn immature INP. Graphs show mean <u>±</u> standard deviation. P-values: ****<0.0001. “ns” indicates no significance.

